# Live-cell micromanipulation of a genomic locus reveals interphase chromatin mechanics

**DOI:** 10.1101/2021.04.20.439763

**Authors:** Veer I. P. Keizer, Simon Grosse-Holz, Maxime Woringer, Laura Zambon, Koceila Aizel, Maud Bongaerts, Lorena Kolar-Znika, Vittore F. Scolari, Sebastian Hoffmann, Edward J. Banigan, Leonid A. Mirny, Maxime Dahan, Daniele Fachinetti, Antoine Coulon

## Abstract

Our understanding of the physical principles organizing the genome in the nucleus is limited by the lack of tools to directly exert and measure forces on interphase chromosomes *in vivo* and probe their material nature. Here, we present a novel approach to actively manipulate a genomic locus using controlled magnetic forces inside the nucleus of a living human cell. We observe viscoelastic displacements over microns within minutes in response to near-picoNewton forces, which are well captured by a Rouse polymer model. Our results highlight the fluidity of chromatin, with a moderate contribution of the surrounding material, revealing the minor role of crosslinks and topological effects and challenging the view that interphase chromatin is a gel-like material. Our new technology opens avenues for future research, from chromosome mechanics to genome functions.

## Introduction

Recent progress in observing and perturbing chromosome conformations has led to an unprecedented understanding of the physical principles at play in shaping the genome in 4D^1^. From genomic loops and topologically associating domains (TADs) to spatially segregated A/B compartments and chromosome territories, the different levels of organization of the eukaryotic genome are thought to arise from various physical phenomena, including phase separation^2–4^, ATP-dependent motors^4,5^, and polymer topological effects^6^. Nonetheless, the physical nature of chromatin and chromosomes inside the nucleus and its functional implications for mechanotransduction remain active areas of investigation^7,8^.

Observation-based studies assessing the mobility of the genome in living cells, from single loci^9–11^ and small regions^12^ to large domains^13^, underline the possible existence of different material states of chromatin (liquid, solid, gel-like). Extra-nuclear mechanical perturbations, including whole-nucleus stretching^14^, micro-pipette aspiration^15^, and application of local pressures^15,16^ or torques^17^ onto a cell, all affect the overall geometry of the nucleus and reveal global viscoelastic properties. Intra-nuclear mechanical manipulation of the genome, on the other hand, is rare and technically challenging^8^. Viscoelasticity measurements using a microinjected 1 μm bead suggested that interphase chromatin may be a crosslinked polymer network (i.e. gel)^18^. Recently, an elegant way to probe intra-nuclear mechanics was demonstrated by monitoring the fusion of both artificial^19^ and naturally-occurring^20^ droplet-like structures. Active mechanical manipulation of an intra-nuclear structure was recently achieved using optical tweezer to displace a whole nucleolus in oocytes^21^ and using optically induced hydrodynamic flows within prophase nuclei^22^. However, all of these approaches are limited to the manipulation of large structures and/or do not apply forces directly on chromatin. These limitations have made it difficult to disentangle various effects (mechanical response of the nucleus *vs*. chromatin itself; hydrodynamics *vs*. polymer viscoelasticity), leading to seemingly contradictory results. Hence, an approach for the direct and active mechanical manipulation of specific genomic loci inside living cells is still needed.

Here, we present a novel technique that achieves targeted micro-manipulation of a specific genomic locus in the nucleus of a living cell, using magnetic nanoparticles (MNPs) and a microfabricated magnetic device. We were able for the first time to exert controlled forces onto a genomic locus and follow its displacement in the nuclear space, giving access to unprecedented measurements on the mechanics of interphase chromosomes in live cells. This new technique opens many avenues for future research, ranging from the biophysics of chromosomes to the perturbation of genome functions.

## Results

### A new approach for *in vivo* mechanical manipulation of a genomic locus

In order to exert a mechanical force onto a chromosomal locus in a living cell, we developed a technique that relies on tethering MNPs onto a genomic repeat and applying an external magnetic field (Fig. 1A). Choosing ferritin for its small size^23,24^ (12nm in diameter; PDB 1GWG), we produced MNPs by synthetizing *in vitro* recombinant eGFP-labeled ferritin cages, loaded with a magnetic core (see *Methods*). MNPs were then microinjected into the nuclei of living human U-2 OS cells harboring an artificial array at a single genomic location (subtelomeric region of chromosome 1; band 1p36) with around 19,000 *tetO* binding sites^25^. MNPs were targeted to the array using a constitutively expressed fusion protein (TetR, mCherry and anti-GFP nanobody) serving as a tether (Fig 1A). Upon injection, MNPs accumulate at the array, forming a fluorescent focus in both eGFP and mCherry channels (Fig. S1A). Quantification of the fluorescence signals indicates that MNPs are at nanomolar concentrations in the nucleus following injection and accumulate at the genomic locus in the range of hundreds to thousands of MNPs (median 1200 MNPs; see Fig. S1B, Fig. S2, Table S1 and *Methods*). The locus should be regarded as a 4 Mb region^25^ (1.6% of chromosome 1) with small MNPs (~2-3 times^24^ the size of a nucleosome) sparsely decorating chromatin (1 MNP per ~15kb). We observe that the locus typically resides in low to intermediate DNA density regions and is itself relatively condensed (Fig. S3). Cells were imaged on a coverglass with custom-made microfabricated pillars^26,27^, which behave as local magnets only when subjected to an external magnetizing field (Fig. S4). Hence, ON/OFF modulation of the local force field can be achieved while imaging by placing/removing an external magnet on the microscope stage. The shape and orientation of the pillars were chosen to maximize the magnetic field gradient and hence the force. We performed magnetic simulations and experimental measurements (see *Methods*; Fig. S5) to determine the magnitude and orientation of the force applied onto the genomic locus, as a function of the number of MNPs bound to it and its position relative to the magnetic pillar (Fig. 1B). The typical forces applied onto the locus were in the sub-picoNewton (pN) range, occasionally reaching a few pN (Table S1). Importantly, these values are in the range of forces exerted by molecular motors in the nucleus, e.g. comparable to the stalling force of ~0.5 pN for the structural maintenance of chromosomes (SMC) complex condensin^28^ and a few pN for RNA polymerase II^29^ (Pol II).

**Fig. 1.**
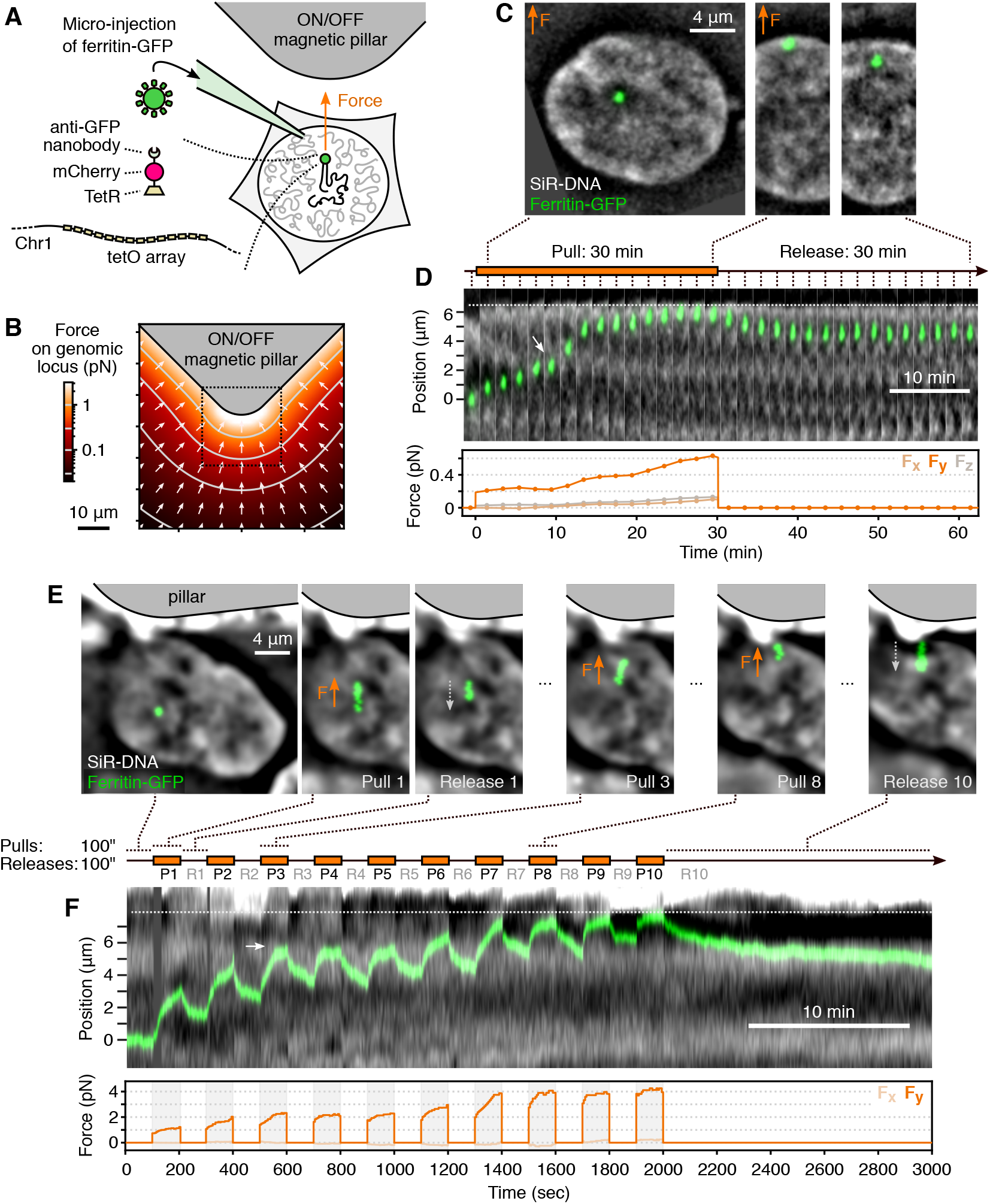
Mechanical micro-manipulation of a genomic locus in living cells. (**A**) Magnetic nanoparticles (MNPs) of GFP-labeled ferritin are microinjected into the cell nucleus and targeted to a genomic array containing ~19,000 tetO binding sites^25^ with a linker protein. Cells are imaged on a coverslide with microfabricated magnetic pillars that produce a local magnetic field and attract the genomic locus. (**B**) The force exerted onto the locus depends on its position relative to the pillar and is characterized using a pre-calculated force map (see *Methods*), here shown for 1000 MNPs at the locus. (**C**) Example of a pull-release experiment showing the locus being displaced during the 30 min of force exertion and recoiling during the 30 min of force release (30’-PR scheme). See also Movie S1. (**D**) Kymograph of the same experiment showing each time frame, along with the force time profile calculated using the force map. (**E**) Experiment where pulls and releases are 100 sec and the pull-release cycle is repeated 10× (100”-PR scheme). Images are time projections, i.e. showing in green all the positions of the center of mass of the locus over the periods represented on the timeline. The arrows indicate the direction of the motion. See also Movie S2. (**F**) Kymograph of the same experiment, showing the displacement of the locus over the DNA density landscape, along with the force time profile. All SiR-DNA images are band-passed (see *Methods*). On (D,F), dotted lines: nuclear periphery, white arrows: feature of interest in the DNA density landscape.

### Force-induced movements of a genomic locus reveals viscoelastic properties of chromatin

We first applied the magnetic force for 30min and released it for another 30min, while performing low-illumination 3D imaging with a Δt=2min interval (30’-PR scheme). We observed a clear motion of the locus toward the magnet upon application of the force and a slow and partial recoil when the force stops (Fig. 1C-D, Movie S1). This indicates that a sub-pN force, when applied in a sustained and unidirectional manner on a genomic locus, elicits a displacement of microns over minutes. It also shows that the locus can move across the nuclear environment, which is believed to be crowded and entangled. We also applied the force periodically–pulling for 100sec, releasing for 100sec and repeating this cycle 10 times (100”-PR scheme)–while performing fast 2D imaging with a Δt=5sec interval (Fig. 1E-F, Movie S2). Several immediate observations from these two experiments already hint at the material properties of chromatin. First, the trajectories show recoils during release periods and a gradual slow-down during both pulls and releases, characteristic of a viscoelastic material. Second, spatial heterogeneities in the trajectories are visible and appear to relate to the spatial DNA density landscape (Fig. 1D and F, white arrows, where locus motion is hindered where the DNA density varies). Third, recoil after force release is seen even after collision with the nuclear periphery, indicating that the material there (peripheral heterochromatin, nuclear lamina) are not sticky enough to retain the locus. Fourth, the DNA density landscape in the nucleus does not show large-scale deformations, excluding that a large amount of material (e.g. the chromosome territory) is dragged along (Movie S1 and Movie S2). Together, our force-induced displacements of a chromosomal locus draw a picture of viscoelastic and non-confining chromatin and provide a rich ground for developing and testing physical models of interphase chromosomes.

### Quantitative force-response and scaling laws of interphase chromatin mechanics

Comparing the trajectories obtained on different cells further highlights and quantifies the viscoelastic properties of chromatin. Analyzing trajectories for the 30’-PR scheme (corrected for cell motion and force orientation, see *Methods*), we observe a range of behaviors for both pulls and releases, regarding initial speed, total distance travelled, and shape of time profiles (Fig. 2A-B). For comparison, we also plotted the envelope of the 100”-PR trajectory (Fig. 2A, green area) and its last release (R10 on Fig. 2B). Most traces show a displacement that is clearly distinguishable from diffusion (Fig. 2A-B, gray areas; see *Methods*). Collision with the nuclear periphery only happens in a minority of cases (Fig. 2A, open symbols) and hence does not account for the range of observed behavior. The initial force applied onto the locus, on the other hand, predicts part of the variability seen in the initial motion (Fig. 2C and Fig. S6A). Similarly, the recoil motion after force release is in large part predicted by the total distance over which the locus has been displaced during the pull (Fig. 2D and Fig. S6B), with a simple linear relationship highlighting the elastic nature of chromatin. Deviations from these simple proportionality relationships indicate how the specific nuclear context of the genomic locus influences its response. In particular, we observe that the two loci that move respectively slower and faster than expected (Fig. 2C, black and gray arrows) are the ones where the locus is respectively the least and the most DNA dense of all analyzed cells (Fig. S3A, arrows), suggesting that the compaction state of the locus itself affects its response to a force.

**Fig. 2.**
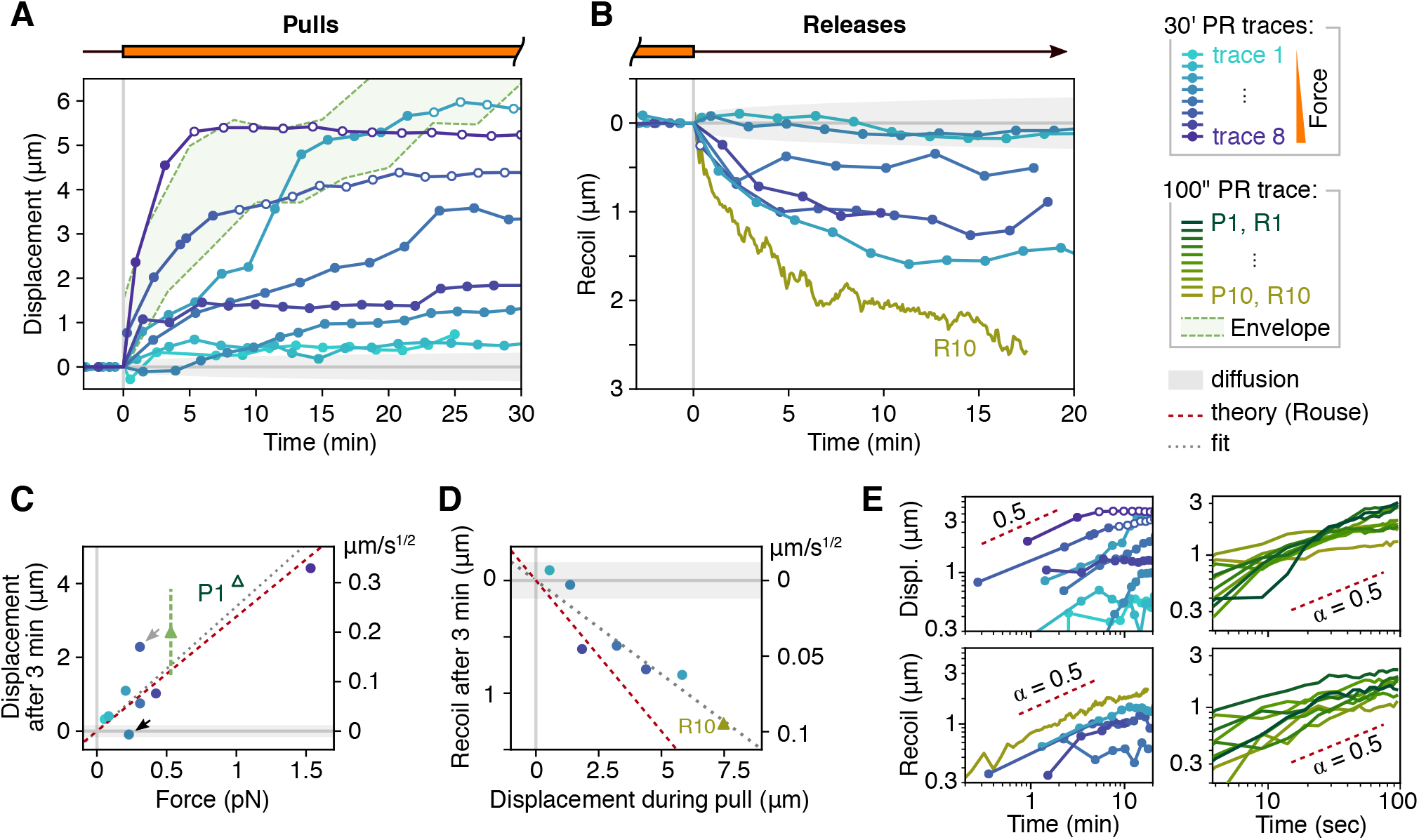
Quantitative analysis of locus movement in response to force. (**A**) Trajectories of the genomic locus in the direction of the applied force for 8 different cells during force exertion with the 30’-PR scheme. Open circles indicate when loci are within 1 μm of the nuclear periphery. Curves are color-coded by force. Green area shows the envelope of the 100”-PR trajectory for comparison. Gray areas in A to D correspond to the null model of pure diffusion based on MSD measurement (see *Methods*). (**B**) Recoil trajectories relative to the time and position at the start of the release are shown for the 6 loci from A where it was recorded. Curve R10 is the last release of the 100”-PR trajectory. (**C**) Displacements measured at Δt=3 min of force exertion on all the traces from A, plotted against the magnitude of the force. Coordinates are interpolated between the frames before/after Δt. The green line/triangle correspond to the 100”-PR envelope shown in A. Reported forces are the average over Δt. Displacements are also expressed in μm/s^1/2^ (right axis), allowing us to place pull P1 from the 100”-PR trace, measured at Δt=100s. The red line indicates the expected relationship from Rouse theory, solely based on an MSD measurement. Arrows indicate specific trajectories deviating from the expected behavior. (**D**) Recoil after Δt=3 min of force release on all the traces from B and the last release of the 100”-PR trajectory (R10), plotted against the total displacement during the pull. The red line indicates the expected relationship from Rouse theory. (**E**) Displacement and recoil trajectories from A and B (left) and from each pull and release of the 100”-PR trajectory (right), aligned on the time and position at the moment of force switching, represented as log-log plots. Straight portions of the curves indicate a power-law behavior, which match the exponent 0.5 predicted by Rouse theory (red lines).

Interestingly, in spite of the variability between traces, log-log plots of all the pulls and releases from the 30’-PR and 100”-PR trajectories together reveal a power-law behavior (i.e. linear portions in the curves), with a slope of 0.5, over two orders of magnitude in time (Fig. 2E). This scale-invariant behavior suggests that the different levels of the hierarchical genome organization are surprisingly not characterized by vastly distinct mechanical properties. In addition, displacements that scale with time as *t*^1/2^ can be empirically described by a ‘fractional speed’, i.e. a single value in μm/s^1/2^ capturing how the motion evolves over time (Fig. 2C-D and Fig. S6A-B, right axes). The first pull of the 100”-PR trace, represented in this unit, indeed follows the same relationships (P1 on Fig. 2C) and the slope of the resulting force-displacement plot yields a unique factor of 0.252 (± 0.026) μm/s^1/2^/pN, characterizing the dynamic response of chromatin to force. Together, these results indicate that a large part of the response of chromatin to force can be described by simple laws.

### Chromatin response to force is well described by a free polymer model (Rouse chain)

We then sought a model of chromatin that best explain our quantitative measurement of force-induced locus displacement. Several features in our data suggest a classical polymer model known as a *Rouse* polymer^30^ as a first approximation to describe the response of chromatin to forces. The Rouse model represents a polymer in which each monomer diffuses by thermal motion in a viscous medium and is attached to its two neighbors by elastic bonds. Importantly, Rouse ignores steric effects (contact, hindrance), crosslinks (affinity, stickiness), and topological effects (fibers can pass through each other). This model is frequently invoked for chromatin dynamics since it predicts the characteristic power-law scaling of the mean square displacements (MSD) *vs*. time, with exponent 0.5, as observed here (Fig. S7A-B) and for other genomic loci in eukaryotes^31–33^. We extended Rouse theory to study how a polymer responds to a point force (see *Methods* and Suppl. text S1). Our calculations predict a power-law behavior with exponent 0.5 for displacements and recoils in response to force, as we observed experimentally (Fig. 2E). These two power laws have the same physical origin, so the diffusion coefficient obtained independently from the MSD (1,979 ±42 nm^2^.s^−1/2^, Fig. S7B) directly relates to –and predicts– the slope of the force-displacement plot (Fig. 2C, red line), i.e. 1,979 nm^2^.s^−1/2^ / 2*k_B_T* = 0.231 ±0.005 μm/s^1/2^/pN; see *Methods*. This agreement between two independent passive and active measurements (diffusion and force response, i.e. red *vs*. gray lines on Fig. 2C and Fig. S6A) strongly supports the Rouse model for chromatin dynamics. After force release, the Rouse model also predicts a recoil proportional to the total displacement during the pull, which we observe qualitatively (Fig. 2D and Fig. S6B) although recoils are somewhat slower than predicted by the theory, suggesting additional effects from the nuclear environment.

### Model-based trajectory analysis reveals moderate hindrance by surrounding chromatin

To further understand the physical nature of chromatin, we asked how alternative polymer models are able to capture the 100”-PR trace (A-D). The approach is to use the displacement trajectory and infer, assuming a given polymer model, the time profile of the force that produced the measured trajectory (see *Methods*, Suppl. text S1 and Fig. S8A-C). Disagreement between predicted and actual force profiles indicates when models are incorrect or incomplete, allowing one to select and refine the best model(s). With this approach, we compared a series of models (Fig. S8C). First, a simple Rouse model without any adjustable parameters (i.e. calibrated using the MSD *vs*. time plot; Fig. S7B) predicts well the first pull and all the release periods (B). However, the prediction leaves some of the applied force unexplained (gray area between curves, B), suggesting a missing ingredient in the model that slows down or hinders the progression of the locus. This residual unexplained force does not scale with speed and hence cannot correspond to extra bulk viscosity of the locus (Fig. S8D). Instead, it increases progressively across successive pulls, suggesting an accumulation of hindrance as the locus progresses through the nucleus. To represent this, we added the capacity for the locus to interact with the surrounding chromatin, represented as extra Rouse chains that are either attached to or pushed by the locus along its path (C and Fig. S8C). These models better predict the force profile throughout the trajectory compared to a pure Rouse model. The only free parameter in C is the frequency at which the locus interacts with other polymers and is strikingly low (Fig. S8C), indicating that the interaction with the surrounding chromatin is moderate. Taken together, these modeling results suggest that chromatin is well described as a Rouse polymer –i.e. a free polymer in a viscous environment– with moderate interactions from the surrounding chromatin, indicating that hindrance, crosslinks, and topological effects play a minor role.

**Fig. 3.**
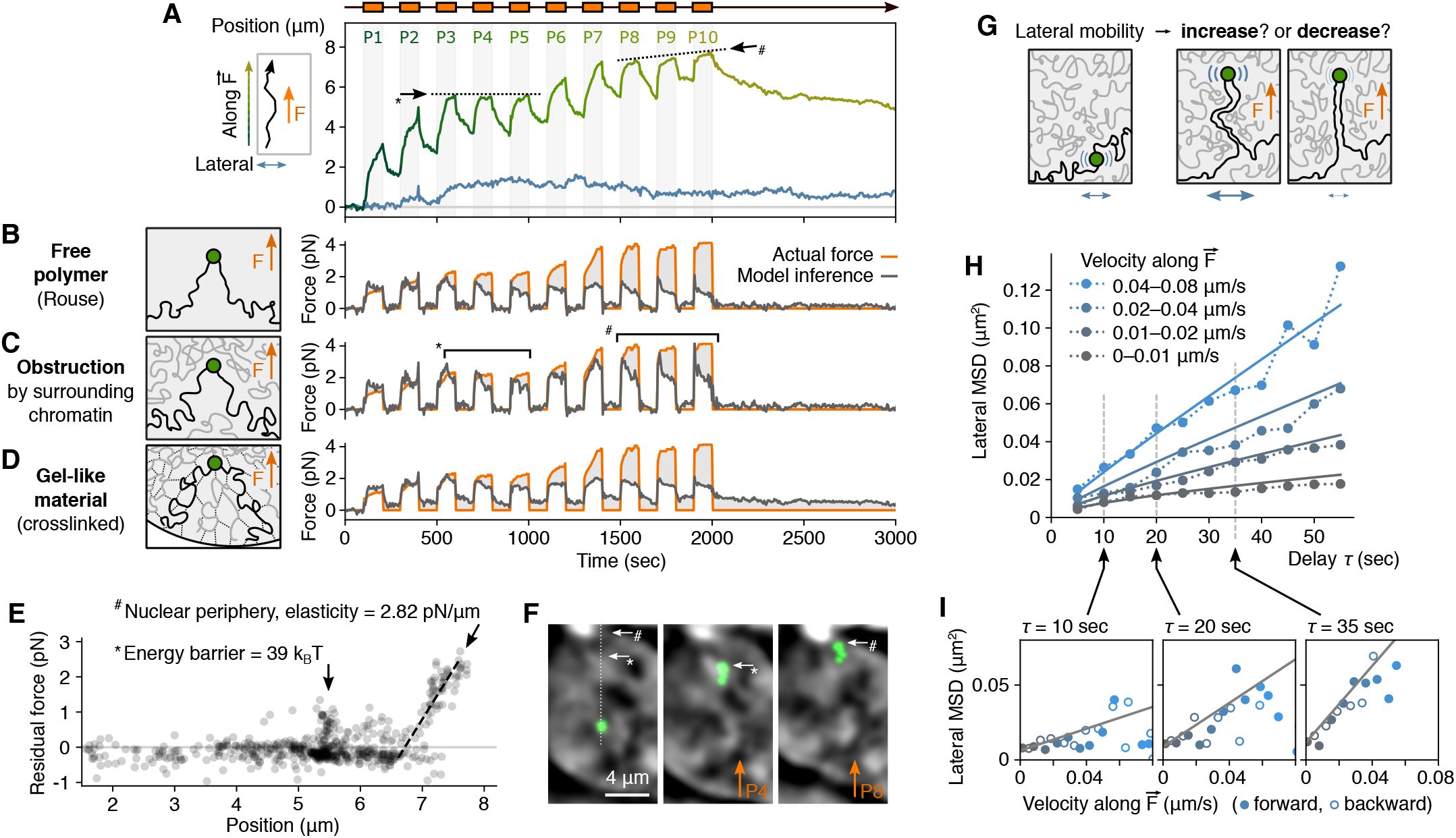
Model-based analysis and hypothesis testing. (**A**) Trajectory of the locus shown in the direction of the force (green curve) and orthogonal to the force in the imaging plane (blue curve) for the of the 100”-PR experiment. Arrows indicate apparent obstacle (see also * and # in panels C, E and F). (**B-D**) Evaluation of different models in their capacity to reproduce the experimentally measured force time profile (orange curve) by inferring it from the trajectory (gray curve). Models shown here are (**B**) a simple Rouse polymer^30^, (**C**) the same model with extra polymer chains being pushed by the locus to represents the surrounding chromatin, and (**D**) a gel-like material, represented as a Rouse polymer in a viscoelastic environment. See full list in Fig. S8. (**E**) The residual unexplained force from the second model (area between curves in C) is plotted along the trajectory of the locus, highlighting an obstacle (*) and an elastic region near the nuclear periphery (#) for which physical parameters are measured (see Methods) and which are visible in panels A, C and F. (**F**) Time projection images, respectively before the first pull and during pulls P4 and P8, showing how the DNA density landscape relates to the identified obstacles. SiR-DNA images are band-passed (see *Methods*). (**G**) Hypotheses on how the lateral mobility of the locus may change depending on its force-induced displacement. (**H-I**) Mean square displacement (MSD) of the lateral movement of the locus, calculated as a function of both time delay and velocity in the direction of the force. Solid lines on both the MSD-delay (H) and the MSD-velocity (I) representations correspond to a single-parameter fit describing how lateral mobility increase with velocity in the direction of the force.

### Interphase chromatin does not behave as a gel in force-response experiments

Chromatin is often proposed to be a gel-like material^11,12,18^. A gel is a highly crosslinked polymer, i.e. many monomers are linked to more than their two neighbors, effectively forming an interconnected mesh with solid-like properties. For chromatin, this could in principle arise from affinity between nucleosomes, as well as loops/bridges formed by proteins/complexes/condensates and topological entanglement between chromatin fibers. First, in such an interconnected mesh structure, short paths effectively linking the pulled locus to all other loci in the nucleus would result in long-range deformation of the DNA landscape, which we do not observe (Fig. 1D,F, Movie S1 and Movie S2). Second, if the chromatin surrounding the locus were gel-like, it would effectively act as a viscoelastic medium. This assumption does not recapitulate well the experimental data (even with two free parameters, D and Fig. S8C) and is inconsistent with the observed scaling of 0.5 in the MSD (Fig. S7B), which argues for a simply viscous and non-elastic medium. Finally, if the locus was part of an interconnected mesh, short series of links would effectively tether it to large structures (e.g. periphery, nucleoli). A Rouse model that includes a finite tether does not recapitulate the experimental data (Fig. S8C) and is inconsistent with the scale-free behavior observed up to several microns (Fig. 2E). These results underline again the minor effects of crosslinks and topological constraints and argue against the view that chromatin behaves like a gel at the spatial and temporal scale of our observations.

### Heterogeneities in the trajectory reveal obstacles in the nuclear interior and a soft elastic material at the nuclear periphery

Even the models that best capture the data leave part of the force unexplained (C, gray area). We thus plotted this residual unexplained force as a function of spatial position (E). This revealed salient features, which match apparent features in the trajectory and the nuclear landscape. First, the residual force in pulls P3 to P5 corresponds to an apparent obstacle in the trajectory (* on A,C) occurring at a high-to-low transition of DNA density (F and Fig. 1F). It appears as a spatially-defined barrier of residual force (E), requiring an energy of ≈39 k_B_T to overcome. This indicates that DNA dense regions are not obstacles *per se*, but rather suggests that the interface between high- and low-density regions may constitute a barrier. The energy we measured indicates that such barriers may be overcome by ATP-dependent molecular motors^28,29^, but unlikely by spontaneous thermal fluctuations. Second, the residual force in pulls P8 to P10 (# on A,C) corresponds to the collision with structures near the nuclear periphery (Fig. 1F and # on F). The observed linear force-distance relationship (E) indicates a solid-like elastic behavior, over at least 800nm and with a spring constant of 2.82 pN/μm. This is surprisingly much softer than the elasticity measured by whole-nucleus stretching experiments^14^, which could be explained by the small size of the locus and/or the existence of a soft layer of elastic peripheral components (heterochromatin, nuclear lamina) before the one contributing to the structural rigidity of the nucleus.

### Lateral mobility of the locus reflects transient collisions with obstacles in the nucleoplasm

To investigate the material encountered by the locus, we further analyzed the lateral motion of the locus as it is pulled and released (A, blue curve). We hypothesized that collisions with obstacles would increase lateral mobility, while the locus being dragged into a more constraining and entangled environment would reduce its mobility (G). After computing the MSD of the lateral motion as a function of both time delay *υ* and velocity *υ_y_* along the direction of the force (H-I and Fig. S7C-D), we observed a clear increase of lateral mobility when the locus moves (both forward and backward; I), suggesting the existence of obstacles that deflect the motion. This additional mobility in the MSD is captured by a term proportional to *υ_y_*, as expected for collisions, and proportional to *τ* (not *τ*^0.5^), as expected if the force due to the obstacles persists in the same direction across several frames, indicating the existence of large obstacles. Indeed, in P3 for instance, the lateral motion clearly shows a directional behavior (Fig. 1E and A). However, our observation holds even when excluding all the timepoints before P4 (Fig. S7E-F), indicating that the collision with obstacles is widespread throughout the nucleus. These results, together with our observation that very few chromatin fibers appear to be carried along with the locus, indicate that obstacles are frequently encountered by the locus, but most interactions are weak and transient.

## Discussion

Our novel approach to micromanipulate a genomic locus inside the nucleus of a living cell provides the first-ever measurements of how an interphase chromosome responds to a point force. Overall, we find that interphase chromatin is surprisingly fluid and behaves like a free polymer, with only a weak obstructive effect of the surrounding chromatin and nucleoplasmic material. Indeed, obstacles are observed along the trajectories, but appear to rarely engage in long-lived interactions with the translocating locus, suggesting a minor role for crosslinks and topological effects in the nucleus. Our work also gives unprecedented access to physical parameters and reveals fundamental scaling laws to describe chromatin mechanics, hence providing a foundation for future theories of genome organization.

Our results appear to contrast with previous experiments depicting chromatin as a stiff, crosslinked polymer gel^11,12,18^. Although genomic loci may generally be spatially constrained^12^, we find that ~pN forces can easily move a compact genomic locus across the nucleus over minute timescales (Fig. 1D,F). This result contrasts with a previous study reporting confined sub-micron displacements over seconds upon application of 65 to 110 pN forces to a 1-μm bead^18^. We propose that our results may be reconciled with previous experiments in one or more of three ways. First, unlike a micron-size bead, the locus in our experiments may be small and deformable enough to pass through the surrounding chromatin. Second, chromatin may contain many, small, gel-like patches, embedded in a bath with liquid, Rouse-like properties. This is also in line with our observation that the transiting locus frequently encounters obstacles. Third, chromatin may be a *weak gel*, i.e. with short-lived crosslinks^11^. Such a gel could continuously maintain a stiff, globally connected network that resists stresses over large length scales, while permitting, fluid-like motions at smaller scales. Future experiments modifying the properties of chromatin and nuclear proteins could clarify how observed micro- and mesoscale mechanics can be reconciled.

Organization of chromosomes that allows movement of genomic loci across large distances under weak forces could have implications for a range of genome functions. Large-scale movements of chromosomes occur during nuclear inversion in rod cell differentiation for nocturnal mammals^34^. Long-range directional motion of specific genes has been reported upon transcriptional activation^35,36^. Giant loops (~5um) were observed to be formed by long and highly transcribed genes and attributed to chromosome fiber stiffening^37^. Large scale, nuclear F-actin dependent relocation of certain double strand break sites to the nuclear periphery was also observed^38^. These DNA-based biological processes require a nuclear organization in which such movements are possible. Our results reveal mechanical properties of chromatin where such large-scale movements would only require weak (near pN) forces. Importantly, although sustained unidirectional forces are unlikely to occur naturally in the nucleus, the magnitude of the forces and the timescale of force exertion in our experiments are comparable to those of molecular motors like SMC complexes and Pol II –i.e. in the sub-pN^28^or low-pN^29^ ranges and applied over minutes (e.g. 10 min for Pol II to elongate through a 25kb gene, 5-30 min for SMC complexes). Thus, molecular motors in the nucleus typically operate in a force range that is sufficient to substantially reorganize the genome in space.

Our new approach opens many avenues for future research, from the study of interphase chromosome mechanics to the perturbation of genome functions, including transcription, replication, DNA damage repair and chromosome segregation.

## Materials and Methods

### Generation of stable cell line

We derived our cell line from the U-2 OS 2-6-3 cell line^25^ harboring a repetitive array of ~200 copies of a construct with 96 copies of the *tetO* binding sequence at a single genomic location in chromosome 1p36. From these cells, we created a stable cell line expressing the TetR-mCherry-antiGFPnanobody plasmid construct through transient transfection and subsequent selection using a selection marker present on the plasmid. Transient transfection was performed using X-tremeGENE HP (Sigma-Aldrich) according to the manufacturers protocol. Cells that contained the plasmid were selected based on their expression of a neo gene using geneticin (Sigma-Aldrich). The optimal dose of geneticin for the selection of cells was found to be 800 μg/ml based on a kill curve experiment (range of concentrations tested: 100 μg/ml – 1000 μg/ml). Single cell clones with low protein expression levels of the TetR construct were selected and amplified.

### Cell culture

Cells were cultured using DMEM/F12, HEPES, no phenol red (Thermofisher), supplemented with 10% FBS (Sigma-Aldrich) and 1% penicillin/streptomycin (Sigma-Aldrich) under conditions of 37 °C and 5% CO_2_. Cells were split regularly using TrypLE (Gibco) to detach cells from the bottom of T-75 or T-25 flaks in which they were cultured.

### Cell adhesion

On the day of the experiment, the magnetic microarray was coated with fibronectin (10 μg/mL in PBS, FC010 Sigma-Aldrich) for 45 minutes at room temperature. Cells were detached using TrypLE (Gibco) from culture flasks and placed to adhere onto the magnetic microarray for at least 4 hours at a concentration of 3×10^5^ cells/array.

### Nuclear staining

One hour prior to the experiment, the media on cells in the magnetic microarray was replaced with culture media containing 1μM SiR-DNA and 10μM verapamil (Spirochrome). Subsequently, media was replaced with normal culture media.

### DNA plasmid

A TetR-mCherry-antiGFPnanobody plasmid was constructed based on a pIRES-CMV-LS-dCas9-mCherry-antiGFPnanobody plasmid (Dahan lab) and a TetR-GFP plasmid (Dahan lab) through amplifications by Q5^®^ High-Fidelity DNA polymerase (NEB, Cat. No M0492S) (primers: CCACCGGTCGCCACCATGGTGAGCAAGGGCGAGGAGGATAACATG, GCCCTTGCTCACCATGGTGGCGACCGGTGGATCCCGGGCCCGCGG, GTCTCCTCTCCGGACTAAAGCGGCCGCGACTCTAGATCATAATCA, GTCGCGGCCGCTTTAGTCCGGAGAGGAGACGGTGACCTGGGTCCC), followed by Gibson assembly (NEB Cat. No NEB, Cat. No. M5510AA) according to the manufacturer’s instructions.

### Magnetic nanoparticles of ferritin-GFP

Magnetic nanoparticles (MNPs) were produced as described in Liße et al.^24^. Briefly, a fusion between monomeric enhanced GFP (mEGFP) and human heavy chain ferritin (HCF) proteins was expressed and purified *in vitro* by size-exclusion chromatography (26/60 Sephacryl S400 column), yielding ferritin cages decorated with 24 mEGFPs. The particles were then PEGylated with PEG2000 to improve stealth properties and subsequently loaded with a magnetite core (Fe_3_O_4_) doped with 5% cobalt, using an autotitration system (Titration Excellence T5, Mettler-Toledo). Magnetization was measured by Vibrating Sample Magnetometry (VSM), yielding a magnetization at saturation of 0.864 ×10^−16^ and 0.850 ×10^−16^ emu/MNP for the two batches used in this study.

Prior to use, an aliquot of 1.5 μM ferritin MNPs was thawed on ice and spun down at 3000 g for 5 minutes. The supernatant was diluted 1:6.5 and used for microinjection experiments.

### Magnetic microarrays

Microarrays of magnetic pillars on glass coverslips were produced as described in Bongaerts et al.^27^ and Toraille, Aizel et al.^26^, using conventional microfabrication techniques. Briefly, 10μm of permalloy (nickel-iron alloy) were deposited by electroplating on a glass coverslip, over 100×100μm square regions obtained by photolithography and arranged every 270μm on a grid pattern (Fig. S4A-C).

Each microarray coverglass was mounted in a custom-made aluminum holder designed to allow the reproducible placement and removal of an external double-magnet while performing imaging^27^. The double magnet (AIP20X20X20, Magnetiques.fr) was measured to produce a uniform 100mT magnetic field, which saturates the micropillars when placed onto the holder (the micropillars saturate at 50~70 mT and show a very low remanence^26^). The microarray coverglass was mounted in the holder at a ~45° angle so that two opposite corners of the square micropillars are aligned with the external magnetic field.

### Nuclear microinjection

Injection of ferritin MNPs into cells was carried out using the InjectMan 4 and FemtoJet 4i systems (Eppendorf). To this end, a Femtotip needle (Eppendorf) was loaded with 3 μl of ferritin and placed onto the injection arm. Typically, a constant pressure of 70 hPa, an injection pressure of 80 hPa and injection time of 0.2 seconds were used. Cells were injected in the nucleus using axial injection at an injection speed of 300 μm/s. Cells were injected at least 1hr before the start of the timelapse experiment and monitored periodically for viability both before and after the time-lapse acquisitions (Fig. S1A, C and D). Following injection, ferritin MNPs typically accumulate at the locus over a few tens of minutes (Fig. S1B).

### Live cell imaging and experimental procedure during acquisition

Cells were imaged using a custom widefield epifluorescence microscope (Nikon, Eclipse Ti2) with a Spectra-X light source (Lumencor), a 100x/1.49NA oil objective (Nikon, CFI Apochromat), an iXon Life 888 EM-CCD camera (Andor), a HLD117NN stage (Prior), a Nano-ZL100 stage piezo (Mad City Labs) and a microscope enclosure chamber (Okolab) to maintain temperature and humidity. The microscope was controlled by the MicroManager software^39^.

Unless stated otherwise, z-stacks of 21 to 24 planes were taken (Δz=0.21 or 0.3 μm) with 100ms exposures and Spectra-X powers at 5% for the GFP, mCherry and SiR-DNA channels. Illumination powers were chosen to be minimal and blue/UV light was excluded, as to avoid light-induced artefacts in chromatin montion^35^. Pixel size is 0.13 μm (or 0.087 μm when an internal 1.5x magnification lens was used, i.e. cells 1 and 2 on Fig. S3)

In the 30’-PR experimental scheme, individual z-stacks were taken in all 3 fluorescence channels prior to the time-lapse experiment. Then, the external magnet was place onto the microarray holder and a Δt=2min time-lapse acquisition was immediately started for 60 min over 6 to 8 stage positions (i.e. micropillar tips), with only the GFP and SiR-DNA channels. The external magnet was removed 30 min into the time-lapse acquisition. Whenever the external magnet is not on the holder, a counterweight was placed onto the holder to minimize mechanical drifts. However, on occasions where placement or removal of the external magnet led to a loss of focus, the acquisition was stopped, adjusted and resumed (note that all analyses rely the actual timestamp of each image to accommodate for the resulting variable frame intervals – see ‘Image analysis’ section). The computer clock time was noted at each placement and removal of the magnet for precise analysis.

In the 100’-PR experimental scheme, a single z-plane was acquired every 5 section in the GFP and SiR-DNA channels for 3000 sec (600 frames) with the same imaging setting as the 30’-PR scheme. After starting the acquisition, the external magnet was placed and removed every 100 sec (20 frames) 10 times, adjusting the focus while acquiring when necessary. After the 10^th^ magnet removal the acquisition was left to finish for the remaining 1000 sec.

### Image analysis

For each field of view in imaged in the 30’-PR experiments, all the acquisitions (i.e. the time-lapse acquisitions, as well as the different single z-stacks taken prior to it) were concatenated into a single movie and individual timestamps were extracted for each frame. 8 cells in total were chosen from 5 different days of acquisition, chosen to represent a range of distance from the micropillar. Specifically, the choice of the cells was based on: (i) proximity to the magnetic micropillar (as to include cells with large forces), (ii) quality of imaging (avoiding loss of focus of the locus, cells close to/overlapping with each other, bright stains on the coverglass) and (iii) apparent cell health (excluding cells that were damaged due to microinjection). For each one of the selected concatenated movies, band-passed versions of the GFP and SiR-DNA channels were generated using Fiji (σ_1_ = 0.8px, σ_2_ = 10px) and treated alongside the raw channels in all subsequent steps. Correction of mechanical drifts was performed using the ‘Correct 3D drift’ Fiji plugin using the micropillar and/or debris on the coverglass as cues. The single-MNP 3D force map (see below) was aligned with the micropillar, merged with the microscopy movie and multiplied by the number of MNPs deduced from the fluorescence intensity of the locus (see below). The resulting movie was rotated to align the force vertically, and the *F_x_* and *F_y_* channels of the force map were adjusted accordingly. The motion and deformation of the cell nucleus was corrected solely based on the SiR-DNA channel, first using the ‘Correc 3D drift’ Fiji plugin and then refined manually using visual landmarks in the area of interest in the nucleus, visualized both on top and side views. 3D tracking of the genomic locus was performed in the resulting movie using a custom-written ImageJ/Python software and the *F_x_*, *F_y_* and *F_z_* components of the force were extracted as the values of the force map at the location of the genomic locus on each frame.

The 100”-PR movie was analyzed in a similar manner as the 30’-PR movie, with few differences: All the steps were performed in 2D since the 100”-PR movie is a single-plane acquisition. The SiR-DNA channel was band-passed using σ_1_ = 2.5px (to reduce further imaging noise) and σ_2_ = 10px. To account for the occasional loss of focus of the locus on a few frames due to the placement/removal of the external magnet, we created an extra channel with a Gaussian at the center of mass of the locus (which hence locates precisely the locus whether in focus or not). This was used for both calculating the trajectory and for visualizations (Fig. 1E-F, F and Movie S2).

All 9 manually-corrected movies (8 with 30’-PR scheme and 1 with 100’s-PR scheme), including raw and band-passed channels, along with all output datafiles (3D trajectories and force), ImageJ scripts and detailed protocol are available (Table S2).

The precise moment of force exertion and force release were determined for each acquisition by keeping notes of the absolute computer clock time of placement and removal of the external magnet with the timestamp of each acquired frame. The position of the locus at each of these two particular instants (which necessarily happen between two frames) was best approximated by taking the position of the locus on the frame just prior to force exertion and force release, respectively.

### Generation and calibration of force maps

The first step to estimate the force applied onto the genomic locus was to generate a force map that describes the 3D force vector per MNP at all positions around the pillar (Fig. S4D-I). For this, we implemented a custom software for magnetic simulations (MagSim, Table S2) and generated simulated force maps, which we calibrated with experimental measurements (Fig. S5).

The simulation software represents the magnetic pillar as a set of finite elements, under a macrospin approximation (i.e. all dipoles are oriented in the same direction, imposed by the external magnetic field 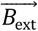). Following classic electromagnetism^40^, the total magnetic field 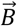 (from all dipoles and 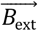) is used to calculate the potential energy 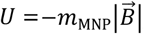, assuming that MNPs are always at saturation (given that our MNPs saturate at ~30 mT, as measured by vibrating-sample magnetometry (VSM)). The 3D vector of the force per MNP is then simply 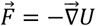. See illustration on Fig. S5A. Besides the geometry of the pillar, the only two parameters are the magnetization of the pillar material *M*_pillar_ and the magnetic moment of MNPs at saturation *m*_MNP_. The latter was measured at 8.5 10^−20^ A.m^2^/MNP by VSM and is reproducible between batches of ferritin MNPs.

To calibrate our simulated force maps, we covered an area of the magnetic microarray with a drop of diluted MNP solution (25 nM) and imaged the spatial distribution of MNPs subject to the magnetic field around the pillars. Using a spinning-disk confocal microscope (Yokogawa CSU-X1 on a Nikon Ti stand with a Plan Apo 100x/1.4 oil objective and an Andor iXon3 camera) to achieve z-sectioning, z-stacks of GFP signal (MNPs) were obtained along with a homogenous control (TMR) in a second channel. Raw z-stack timeseries images were corrected for intensity offset and non-homogenous illumination/light collection (‘flat-field’ correction), time-averaged and down-sampled to minimize noise, and the GFP channel (MNPs) was normalized by the TMR channel (homogenous control) to correct for shadowing effects due to partial obstruction of the confocal light beam at the vicinity of the pillar. Within an intermediate range of fluorescence intensity, the resulting images are assumed to approximate a Boltzmann distribution of MNPs, so that 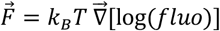. See illustration on Fig. Fig. S5B. For the comparison with the simulated maps, masks were made to exclude areas of low fluorescence signal (prone to noise and inaccurate flat-field correction) and areas of high signal (prone to non-linearity due to steric exclusion and fluorescence quenching). Since even a small amount of photobleaching during the spinning-disk acquisition makes the *z* component of the experimental force maps inaccurate, only the *x* and *y* components were used for comparison with the simulated maps. Raw and processed images and Imagej analysis scripts are available (Table S2).

A single-parameter fit between our simulated map and 6 experimental force maps from 6 different pillars (Fig. S5C-D) yielded a value of *M*_pillar_ = 3.67 10^5^ A/m (which is around the expected value for a saturated pillar of this geometry and composition^26^ and an estimation of the error on the force map below 0.1 fN/MNP. Python library of the MagSim simulation software and Jupyter notebook are available (Table S2).

This calibration was performed for all combinations of microarray coverslips and MNP batches used in this study, and found to be equivalent. A specific map was however produced for each microarray, accounting for the precise orientation of the microarray after it was mounted in the holder. Final force map TIF files are available (Table S2).

### Fluorescence-to-MNPs conversion factor

To deduce the force applied onto the genomic from the single-MNP force maps, we estimated the number of MNPs at the locus in each injected cell using a fluorometric approach. First, we made sure that the accumulation of MNPs within the small volume of the locus did not induced quenching of the GFPs. Indeed, monitoring the redistribution of the GFP fluorescence following injection revealed that, while the MNPs relocate from the freely diffusion pool to the locus, the overall nuclear signal remains constant (Fig. S1B), indicating no major loss of fluorescence when MNPs are at the locus. Second, to convert the measured fluorescence intensity at the locus into a number of MNPs, we estimated the fluorescence of single ferritin MNPs. For this, we used imaging conditions where we can see individual ferritin MNPs, similar to experiments using ferritin or other fluorescent multiprotein nanoparticles as passive tracers^41^. Here, we injected low amounts of MNPs into cells and performed short-exposure (5 ms) max-power imaging (Fig. S2A). After temporal binning of the images and exclusion of bright and static objects, single MNPs were detected and averaged into a single-MNP image (Fig. S2B). The integrated fluorescence intensity was computed and converted into a ‘fluorescence-to-MNP conversion factor’ accounting for our default imaging conditions used for time-lapse acquisitions.

The number of MNPs at the locus varied substantially from cell to cell, typically from a few hundreds to a few thousands (median 1200 MNPs; Table S1). The fraction of MNPs at the locus ranges from a few percent to more than half of the injected pool (median 26%; Table S1).

To calculate the force on the locus, the amount of MNP is calculated on a cell-by-cell basis from the first z-stack of the timelapse acquisition, and is multiplied by the 3D force vector from the single-MNP force map on a frame-by-frame basis (since the locus and the cell move over time relative to the micropillar).

### MSD analysis

Mean square displacements (MSD) were calculated as 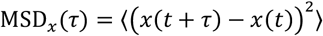, where 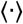 denotes time average. MSD curves were computed on different portions of the 100”-PR trace using the coordinate *x*(*t*) along the axis orthogonal to the force. Computed over the whole 1000 sec of the last release (R10), it clearly indicates a power-law behavior with a slope 0.5, i.e. following MSD_*x*_(*τ*) = Γ *τ*^0.5^ (Fig. S7B). To estimate precisely the pre-factor Γ on this particular cell, we recomputed the same MSD curve only over the last 100 timepoints of the trajectory and fit the first 4 points of the resulting curve, yielding Γ = 1,979 ± 42 nm^2^.s^−1/2^ (Fig. S7A).

The plots of H,I and Fig. S7C-F were made as follows: For all the pairs of timepoints separated by a given delay *τ* in the 100”-PR trace, we calculated the square displacements of the lateral motion SD = (*x*(*t* + *τ*) – *x*(*t*))^2^ and the velocity along the direction of the force *υ_y_* = (*y*(*t* + *τ*) – *y*(*t*))/*τ*. Then, after defining bins for *υ_y_* values (e.g. 0-0.01, 0.01-0.02, 0.02-0.04, 0.04-0.08 μm/s), we calculated the mean of all SD values for which *υ_y_*, fall in the same bin, resulting in an MSD as a function of τ and #$. Re-using the value of Γ from above, a single-parameter fit of the resulting MSD values as 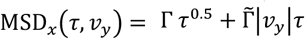 yielded 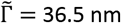.

### Polymer modeling and model comparison

The theory behind the model-based analysis presented on and Fig. S8 is described in depth in Suppl. text S1. We compare here how different polymer models are able to reproduce the observed data: For each assumed model, we calculate the time profile of the force that needs to be applied to the locus in order to produce the observed trajectory. Differences between the force time profile we inferred from the model and the one we measured experimentally indicate if and how the different models capture the observed behavior.

The 6 models we examined are explained on Fig. S8C and formally defined in Suppl. text S1.

Model (i) on Fig. S8C is an infinitely long Rouse polymer, i.e. an approximation that ignores steric and topological effects and where the polymer is not attached to any fixed landmark. This simple model is motivated by the observed power laws with exponent 0.5 in the displacements (Fig. 2E) and the MSD (Fig. S7B). Without any adjusted parameters, this model captures well the first pull and all the recoils –confirming it as a good first approximation, but highlighting its failure to capture the data when the locus is pulled over large distances.

Model (i) predicts a force during the late pulls that is weaker than the actual force. Taken differently, given the actual force profile, the locus should move faster than we observe if the model was accurate and complete. The unexplained force (gray area between curves for model (i) on Fig. S8C) can be viewed as an force acting against the motion of the locus, that is missing in a simple Rouse polymer model. We reasoned that the locus, being decorated with MNPs, could experience a larger viscous drag than the rest of the polymer. However, this is not supported by that fact that the unexplained force does not scale positively with speed (Fig. S8D) and that we do not observe any substantial unexplained force when the locus moves backward during the recoils. Instead, the unexplained force is only seen when the locus moves forward and it increases progressively as the locus is displaced over larger and larger distances. This suggests the presence of material that hinders the progression of the locus and accumulates as it moves across space. Hence, we incorporated an additional force representing the obstruction by the surrounding chromatin, modeled as extra Rouse chains that are encountered and interact with the locus along its path. This is implemented in 3 variants (model (ii) to (vi) on Fig. S8C). In model (ii), the extra chromatin chains stick to the locus permanently and are hence carried along during both pulls and releases. In model (iii) and (iv), the extra chromatin chains are simply pushed without sticking to the locus. Reasoning that less chromatin may be encountered during the release because it has been cleared up during the previous pull, model (iii) include encounters with extra chromatin chains exclusively during forward motion, while model (iv) consider encounters on both forward and backward motion. All three models appear to explain equally well the force profile (Fig. S8C) so that we cannot favor one over the others. However, model (ii), did not perform well if the force-displacement parameter (obtained from Fig. 2C) was not re-adjusted, and even so, it reproduced less precisely the releases periods than models (iii) and (iv). Overall, this highlights the importance of the surrounding chromatin in the dynamics.

In contrast, when we represented properties expected for gel-like materials, the models perform worse. First, we considered a situation where the polymer is embedded in a gel-like medium. Such a medium is known to have viscoelastic properties that would change the way the monomers of our polymer diffuse over time^42^. Model (v) on Fig. S8C, which represents this situation, fails to capture the observed force profile, disqualifying it as an appropriate model. Second, in the case where chromatin would be a gel throughout the nucleus, a series of crosslinks would exist between the locus and fixed landmarks like the nuclear periphery. Model (vi) on Fig. S8C approximates this situation by representing a Rouse polymer where a monomer along the polymer is tethered to a fix position is space. The displacement is hence dominated by the extension and relaxation of this finite tether, which has temporal characteristics that fail to capture our data, even when adjusting for the relaxation time of the finite tether. Indeed, after two pull-release cycles, neither pulls nor recoils are accurately recapitulated.

The force that remains unexplained in models (ii) to (iv) corresponds to visible feature in the trajectory and in the DNA landscape, which we describe as obstacles and interpret quantitatively thanks to our approach (A,C,E,F).

### Theoretical relationship between MSD curve and force-displacement plot

Analytical derivations on the Rouse model indicate that the MSD due to passive diffusion and the displacement in response to force have the form MSD_*x*_(*τ*) = Σ *τ*^0.5^ and 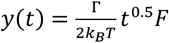 respectively (Suppl. text S1, eqs. (5) and (7)), where *F* is the force applied to the locus and Γ is a constant prefactor of the MSD. Hence, the slope of the force-displacement plot (dimensions [m].[s]^−1/2^.[N]^−1^) is simply the prefactor Γ of the MSD (dimensions [m]^2^.[s]^−1/2^) divided by 2*k_B_T* (dimensions [N].[m]). Experimentally, we measured the slope of the force-displacement plot to be 0.252 ±0.026 μm/s^1/2^/pN (fit on Fig. 2C), which is close to the value of 0.231 ±0.005 μm/s^1/2^/pN predicted from the prefactor 1,979 ±42 nm^2^.s^−1/2^ of the MSD (fit on Fig. S7B).

### Energy and elasticity measurements

From the residual force plot (E), we estimated the energy barrier as the product of the width and height of the cloud of points laying between positions 5.2 μm and 5.7 μm. The elasticity close to the nuclear periphery was computed by fitting a straight line through all the points at positions greater than 6.5 μm.

## Acknowledgments

We dedicate this work to the memory of Maxime Dahan, who died in July 2018. The authors would like to acknowledge scientific inputs from: C. Giovannangeli and J.-P. Concordet (MNHN), T. Pons and N. Lequeux (ESPCI), M. Coppey (Institut Curie), and the members of the Coulon and Fachinetti teams. We acknowledge the current and past members of the Dahan/Coppey/Hajj team and UMR168, including C. Monzel, D. Liße, J. Manzi, E. Baloul, C. Vicario and D. Normanno, for their earlier work and technical help on nanoparticles and microinjection, and E. Secret and A. Michel-Tourgis (UMR8234 CNRS) for help with magnetic nanoparticle characterization. The authors acknowledge the Flow Cytometry Core Facility of the Institut Curie, the BMBC platform of UMR168, the machine shop of UMR168, the technological platform of the Institut Pierre-Gilles de Gennes (IPGG) for access to microfabrication facilities, the PICT-IBiSA@Pasteur Imaging Facility of the Institut Curie (member of the France Bioimaging National Infrastructure; ANR-10-INBS-04), and the MPBT platform (Physical Measurements at Low Temperatures) of Sorbonne Université.

## Author contributions

MD, DF and AC conceptualized the project. MD, DF, VIPK and AC obtained funding. VIPK, MD, DF and AC designed the experiments. VIPK, LZ and AC obtained the data. VIPK, LZ, KA, MB, LKZ, SH produced reagents and provided key technical expertise. MW, VIPK and AC preformed image analysis. MW, VIPK, SGH, VFS and AC analyzed trajectories. SGH, EJB and LAM preformed polymer modeling. VIPK, SGH, MW, VFS, EJB, LAM, DF and AC interpreted the results. DF and AC supervised the experimental work. AC supervised the image analysis work. EJB and LAM supervised the modeling work. AC drafted the manuscript with inputs from VIPK, MW, DF, SGH, EJB and LAM. All authors participated in reviewing and editing the manuscript.

## Data and materials availability

All data and codes are available as detailed in Table S2.

