## Supplementary figures, tables and movie captions for "Live-cell micromanipulation of a genomic locus reveals interphase chromatin mechanics"

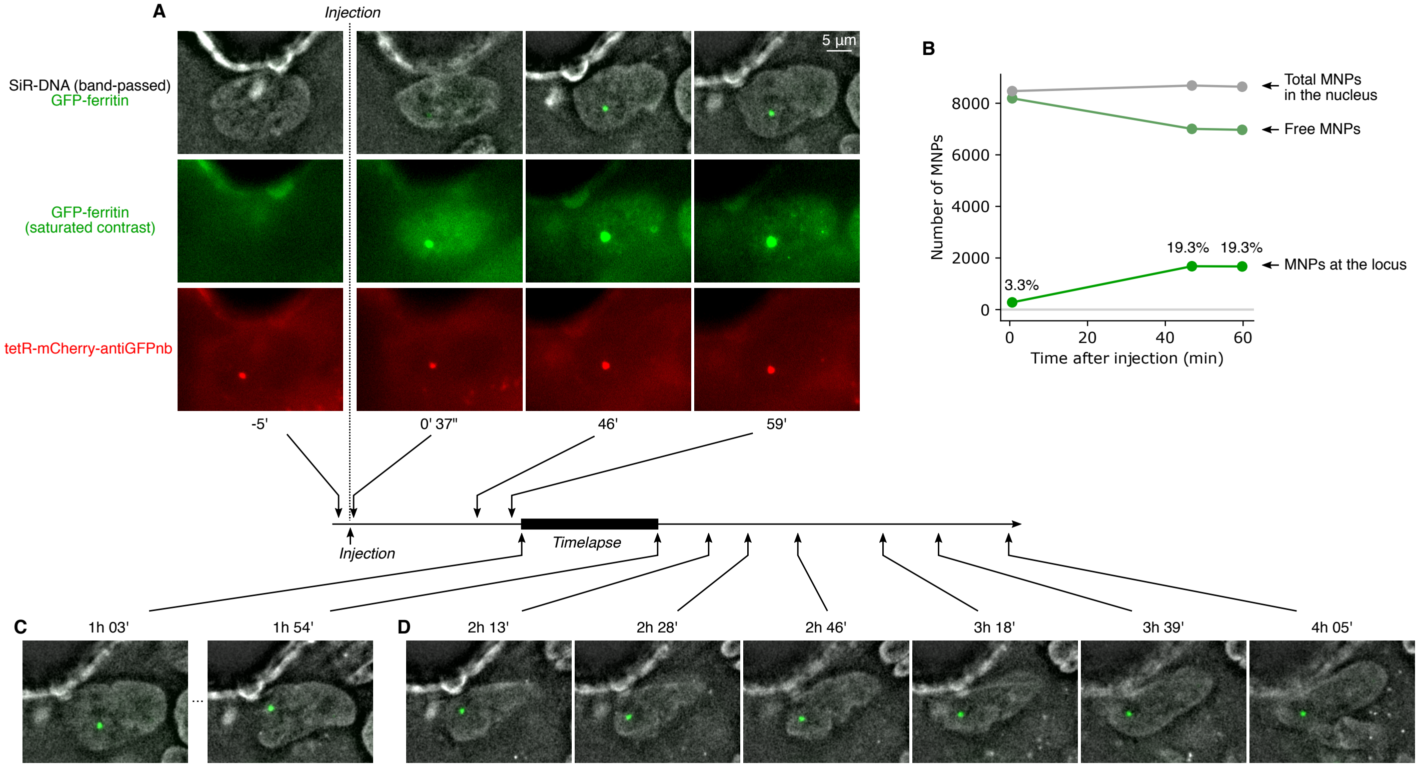


Fig. S1. **Microinjection of MNPs and monitoring of the cell before and after the experiment**. (**A**) As illustrated here with the cell used for the 100"-PR experiment, cells are injected at least 1h prior to the time-lapse experiment. The locus is visible in the mCherry channel before injection and gets progressively loaded with MNPs (First row: GFP channel is not saturated, showing the progressive loading of the locus over time. Second row: GFP channel is saturated in order to visualize the unbound MNPs in the nucleus). (**B**) Quantification of the fluorescence signal at the locus and in the rest of the nucleus. MNPs are observed to rapidly redistribute to the locus and to be fully equilibrated at 45 min post-injection. The total fluorescence in the nucleus remains constant, indicating that fluorescence quenching at the locus is minimal or inexistent. The number of MNPs can hence be determined using a fluorometric approach (see Methods and Fig. S2), yielding an absolute count of MNPs at the locus and in the nucleus. The total amount of injected MNPs here was ~8,000, corresponding to ~10nM (assuming a 15~20µm wide and 5µm thick nucleus). (**C**) The cell was imaged for the micromanipulation experiments for 50 min. (**D**) After the experiment, the cell was monitored for up to 2hrs. All images are taken with minimal illumination powers, sufficient to observe the necessary signals, as to minimize possible side effects of illumination on chromatin dynamics[35]. SiR-DNA channels are band-passed (see *Methods*) and all images are the single z-plane where the locus is in focus.

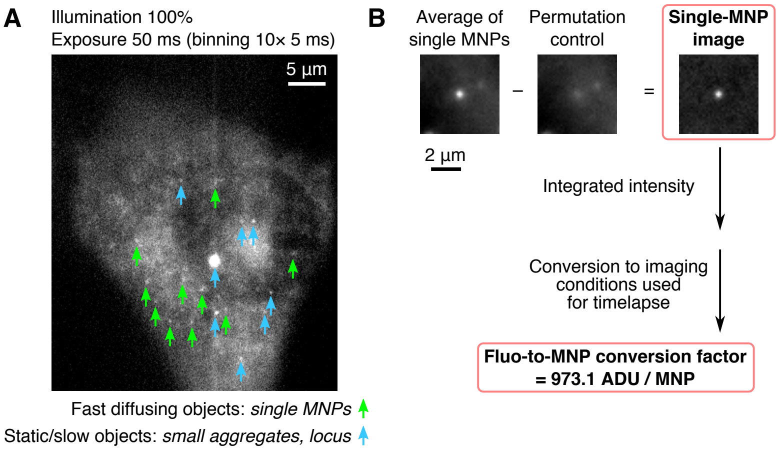


Fig. S2 **Quantification single-MNP fluorescence**. As explained in *Methods*, a 'Fluorescence-to-MNPs' conversion factor was calculated as follows. (**A**) A cell was microinjected with a low concentration of MNPs in both the nucleus and the cytoplasm and imaged with maximal illumination power and 5 ms exposure time in order to visualize single MNPs. Images were subsequently binned 10 by 10, yielding an effective exposure of 50 ms. Diffraction-limited object were located and static or slow-diffusing objects were eliminated (blue arrow) to only retain the fast-diffusing ones (green arrows). (**B**) An average image of all the fast-diffusing particles was computed and subtracted with a control image (made in a similar way, with the same particle positions, but using a later/earlier frame in the time-lapse). The result is a background-free high-quality image of a single MNP. The total fluorescence was measured in a circular area around the MNP and converted into the equivalent value used for our default time-lapse imaging conditions (i.e. 100 ms exposure and an illumination power 7.35x lower, as measured with a powermeter). This value is the conversion factor to estimate the number of MNPs at the locus in our experiments.

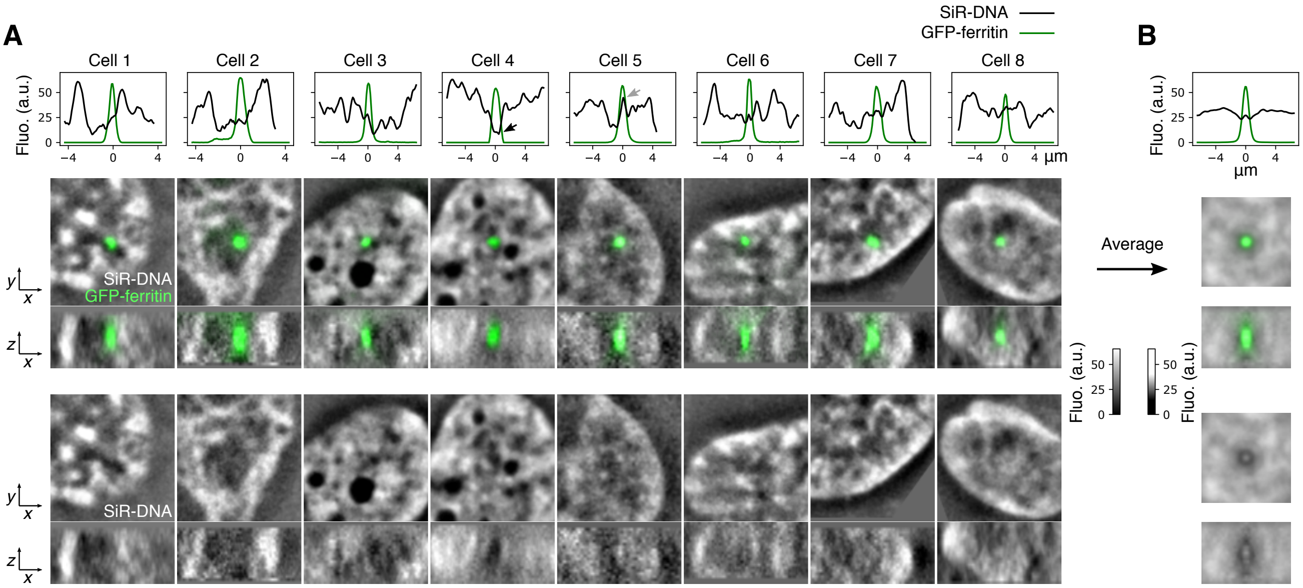


Fig. S3. **DNA density at and around the targeted genomic locus**. (**A**) For each of the 8 cells on Fig. 2 for the 30'-PR scheme, images at the beginning of the experiment are quantified: The line plots (first row) along the *x* direction are shown, indicating that the locus (green peak) often resides in an intermediate DNA density region compared to the neighboring environment. Black and gray arrows identify the same two cells as on Fig. 2C, where the locus shows respectively low and high DNA densities relative to the rest of the nucleus. Top views and side views are shown with both GFP and SiR-DNA channels (middle row) and SiR-DNA channel only (bottom row) to appreciate the density at the locus. Images right before force exertion are shown, except for cell 1, for which the SiR-DNA channel is missing (the first image after force exertion is shown). (**B**) A meta-locus analysis is performed by averaging together all 8 cells and over 10 random rotation angles. The result is represented similarly as panel A. The line plot and the top and side views show that the locus resides in a region which is on average less dense that then mean density in the nucleus and that the locus itself is moderately DNA dense. Note: SiR-DNA images on panel B are shown with a different grayscale as to emphasize subtle variations in density. All SiR-DNA images are band-passed (see *Methods*). Contrast of each image was adjusted to account for the variability in the SiR-DNA staining between experiments. xy views are the average of 5 z-planes centered on the locus.

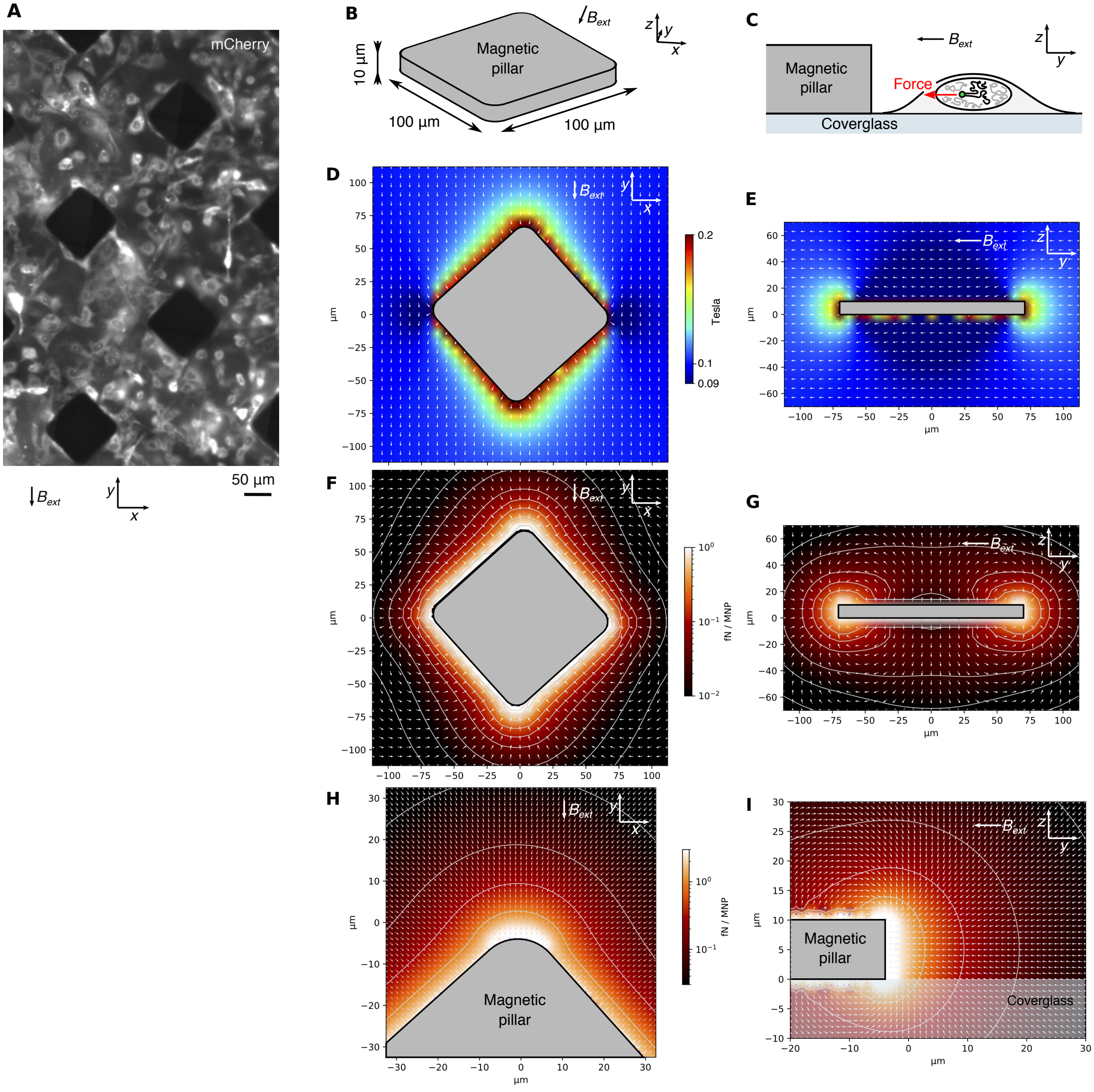


Fig. S4. **Magnetic micropillars and 3D force field**. (**A**) Area of a coverglass showing several microfabricated pillars (large black squares) along with cells expressing the tetR-mCherry-antiGFPnb protein (mCherry fluorescence) Pillars are oriented 45° relative to the external magnetic field $\overrightarrow{B_{\mathrm{ext}}}$, which is applied in the *y* direction. (**B**) Pillars are 100µm×100µm and 10µm in height. (**C**) Side-view schematic of a cell close to the tips of a pillar. Note: the targeted genomic locus is typically found to be 2 to 6µm above the coverglass. (**D**,**E**) Top view and side view of the simulated magnetic field produced by the magnetic micropillar and the external magnet. See Fig. S5 for details on the simulation and its calibration. Top view: z=5µm. (**F**,**G**) Top view and side view of the resulting force field per MNP. (**H**,**I**) Zoomed-in views of F and G, to appreciate the force field close to the tip of the pillar, i.e. where we chose the cells to image as to maximize the force exerted on the locus.

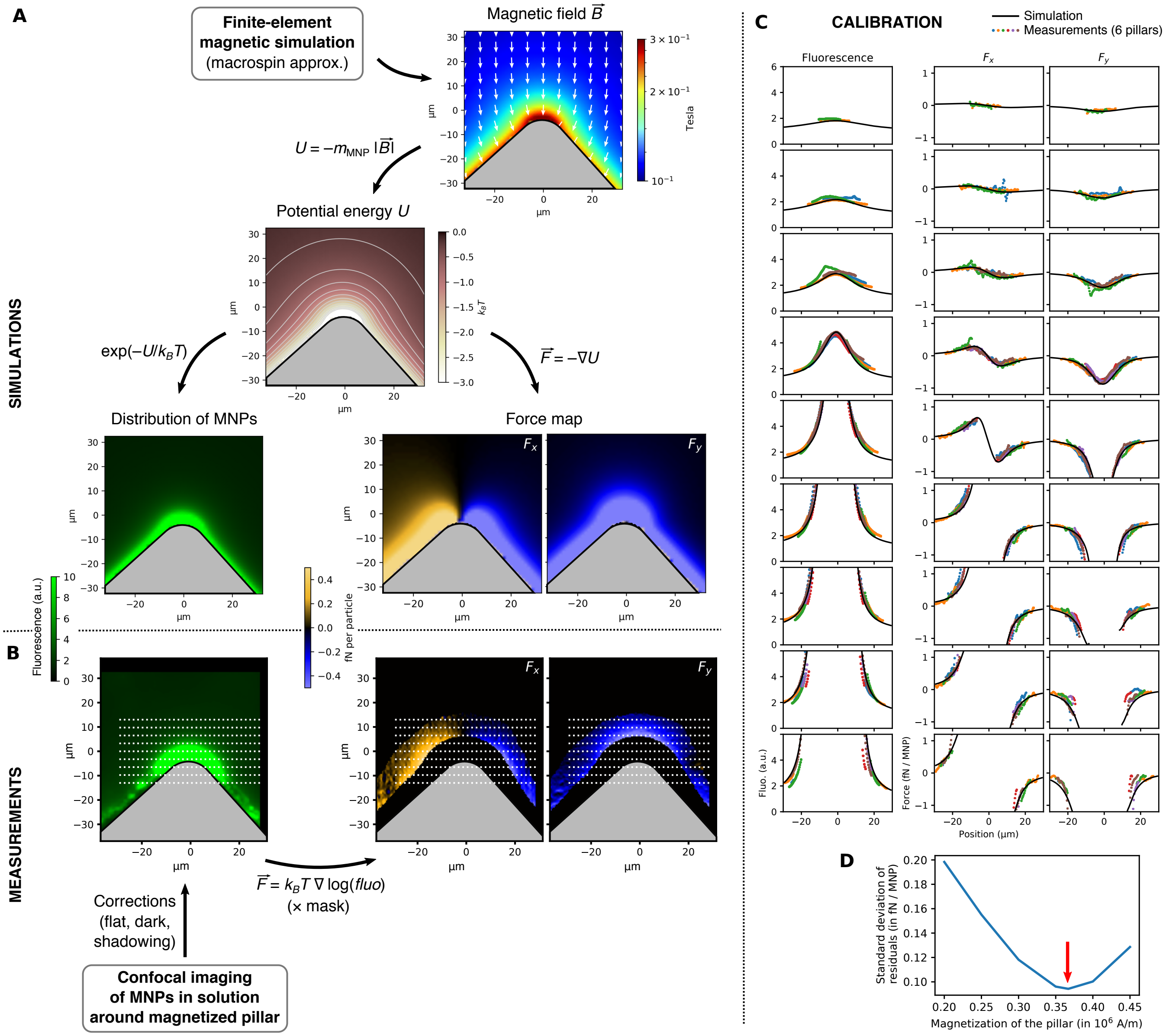


Fig. S5. **Magnetic force map simulation and calibration**. (**A**) Magnetic force simulation was performed using our custom-written software MagSim (Table S2). It uses a finite-element approach to produce a magnetic field, from which we deduce a potential energy field for a single MNP. From this, we deduced both the stationary distribution of particles in space as a Boltzmann distribution and the force in $x$, $y$ and $z$ directions (only $F_x$ and $F_y$ are shown and used for calibration). See *Methods* for details. (**B**) To calibrate the simulations, we performed confocal (i.e. z-sectioned) imaging of MNPs in solution around magnetized pillars (see *Methods* for details). The $F_x$ and $F_y$ components of the force was deduced from the fluorescence within the area where fluorescence is expected to reflect a Boltzmann distribution of MNPs (see *Methods*). (**C**) Simulations (curves) and experimental data from 6 different pillars (colored dots) were compared as series of line plots (dotted lines on B), both in terms of predicted fluorescence and predicted $F_x$ and $F_y$ components of the force. (**D**) The single unknown parameter of the simulation (i.e. the magnetization $M_{\text{pillar}}$ of the pillar material) was determined by minimizing the mean square difference between simulated and measured values of $F_x$ and $F_y$ over all the line plots and the 6 pillars.

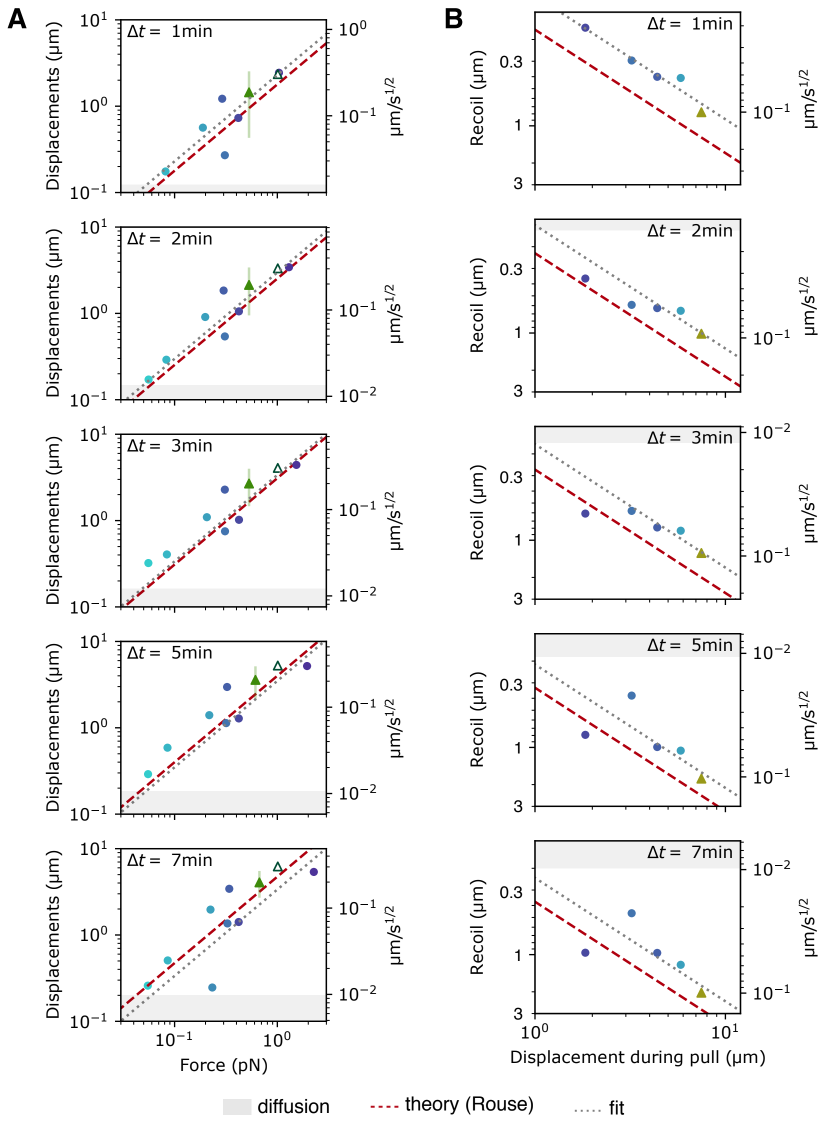


Fig. S6. **Force-response and recoil plots**. (**A**) Scatter plots similar to Fig. 2C and (**B**) Fig. 2D are represented here with log-log axes and for different time delays Δt after the start of the pull (A) or the start of the release (B). Symbol colors and shapes are identical to Fig. 2: Circles are individual trajectories from the 30'-PR scheme, the open triangle represents pull P1 from the 100"-PR experiment, the solid triangle and the vertical line on (A) represents the envelope of the 100"-PR trajectory, and the solid triangle on (B) represents the last release R10 from the 100"-PR experiment. Note that one point from Fig. 2C and one point from Fig. 2D, which have negative values, are not visible on this log-log representation. The gray areas correspond to the null hypothesis of pure diffusion based on MSD measurement (see *Methods*). The gray line is a linear fit to all the data points. The red line indicates the expected relationship from Rouse theory.

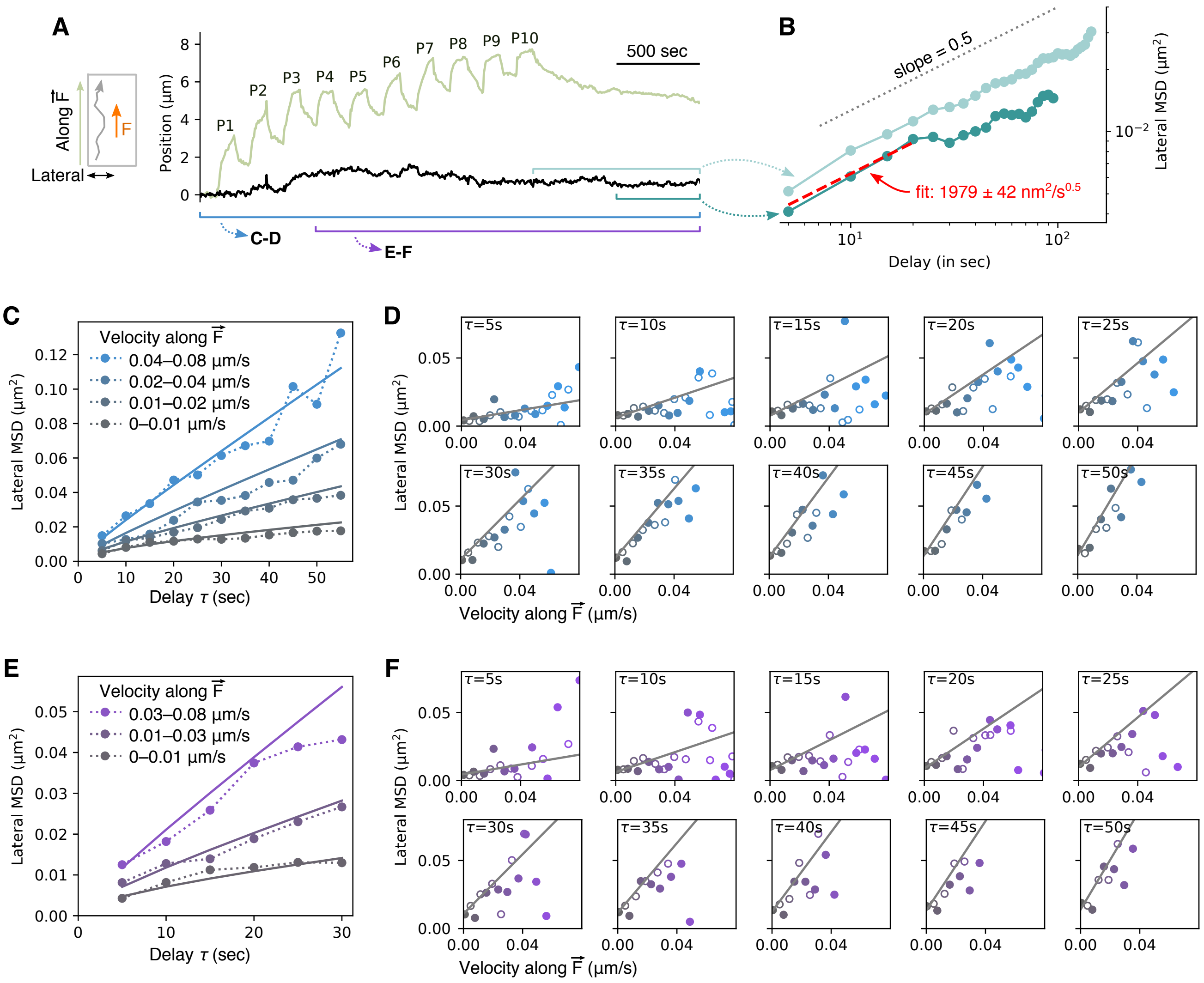

Fig. S7. **MSD and lateral mobility**. (**A**) The lateral component of the trajectory of the locus in the 100”-PR experiment is shown (black curve) and analyzed thereafter in different ways in relation with the trajectory in the direction of the force (green curve). (**B**) A mean square displacement (MSD) calculated on the lateral component of the trajectory over the entire last release period (light curve) shows a power-law behavior with slope 0.5, over nearly two orders of magnitude in time. The MSD calculated over the second half of the last release (dark curve), although less precise since calculated with fewer data points, shows the same power-law behavior, but with a different offset, suggesting that the extent of the mobility depends on the motion of the locus in the direction of the pull-release force. MSD being most accurate at short delays, we fit the latter curve over the first four time-delay points and obtained a prefactor of 1,979 ±42 $nm^2.s^{-1/2}$. (**C**,**D**) The square displacements (SD) over the entire lateral trajectory were calculated for time delays up to 55 sec, then grouped by both time delay and velocity along the force, and finally average within each group, to yield the MSD as a function of both time delay and velocity along the force (see *Methods*). (**E**,**F**) The same analysis as (C,D) was performed only using the trajectory staring at pull P4 (shown on A).

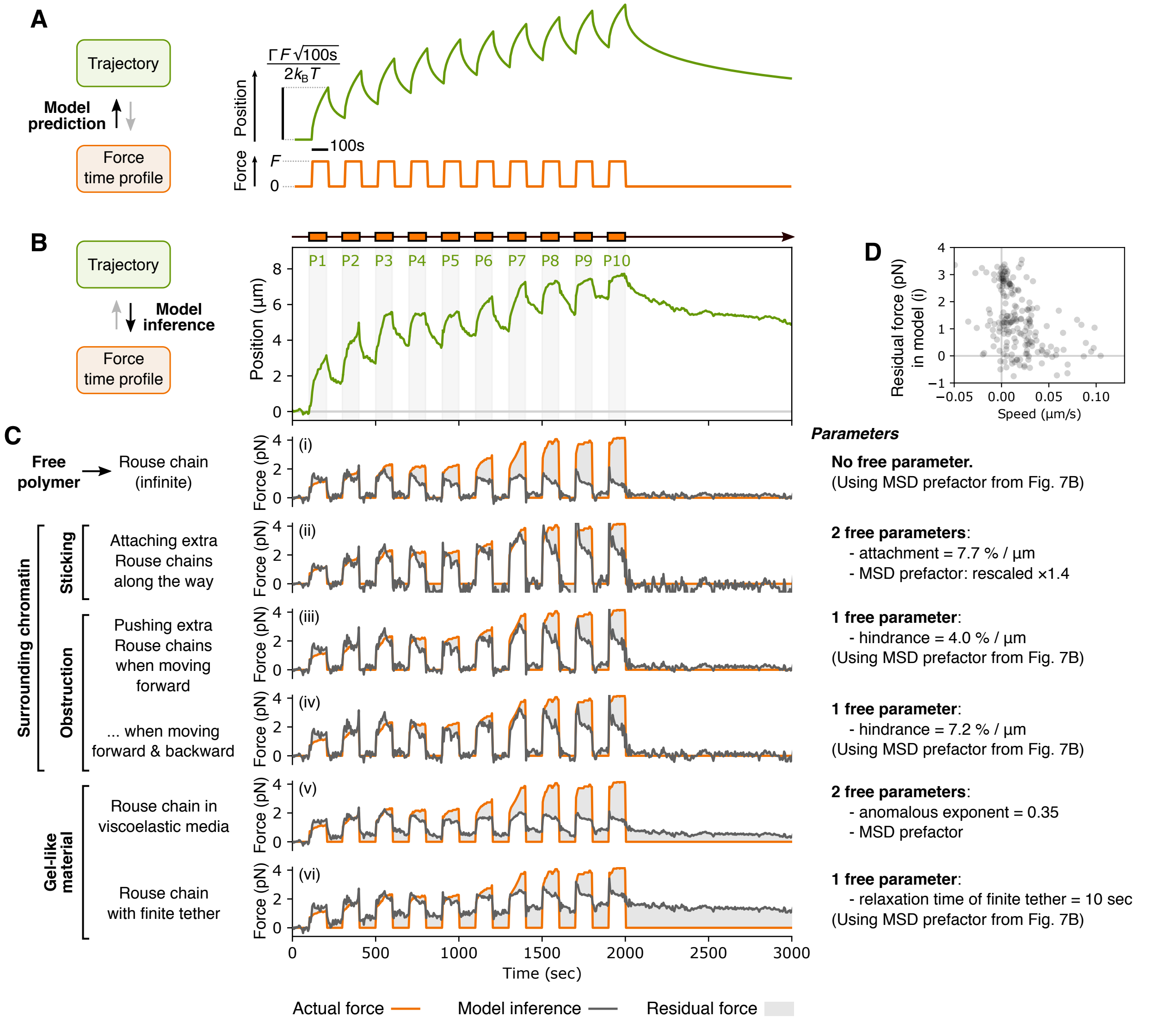


Fig. S8. **Comparison of concurrent physical models using a force-inference approach**. Our modeling approach creates a one-to-one mapping between the time profile of the applied force and the trajectory (Suppl. text S1, section II). (**A**) We can therefore predict the trajectory of the locus for a given force profile (shown is the prediction for a Rouse polymer), or alternatively (**B**) infer the force profile necessary to move the locus along the observed trajectory. We chose inference over prediction, since interpretation of model mismatches (inferred *vs.* measured force) is more straightforward. All models tested below take as an input the trajectory of the locus in the direction of the force from the 100"-PR experiment. (**C**) A series of models are evaluated on their capacity to predict a force time profile (black curves) that recapitulates the measured one (orange curves). The gray area between the curves is the residual force, i.e. the part of the observed force that is not explained by the model. Model (i) is an infinite Rouse polymer, i.e. a free polymer where interactions, hindrance, crosslinks and topological entanglement are ignored (Suppl. text S1, section III). The other 5 models (ii-vi) are variations of this simple model (Suppl. text S1, section V). The first three models consider a polymer as in (i) which, in addition, interacts with the surrounding chromatin, represented as extra polymer chains. These extra chains either (ii) make permanent crosslinks with the locus when encountered, (iii) are pushed by the locus when it moves forward, or (iv) are pushed by the locus when it moves both forward and backward. Leaving open the possibility that the locus is already interacting with extra chains at t=0, the density of extra polymer chains in models (ii-iv) –i.e. the effective number of chains interacting with the locus per µm travelled– is expressed in percent of the initial load. The last two models represent elastic properties that would be expected for a gel-like material: In model (v), we consider that media in which the polymer evolves has itself viscoelastic properties. In model (vi), we represent crosslinks as effectively making the polymer not infinite but attached to a fixed landmark (e.g. nuclear periphery). (**D**) The residual force in model (i) does not scale positively with speed, suggesting that it is not due to extra viscosity of the locus.

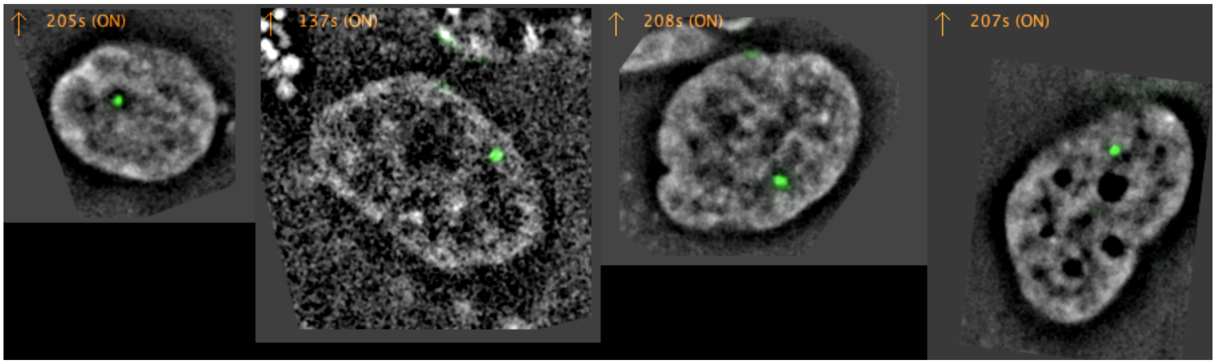


Movie S1. **Examples of 30 min pull-release experiments**. Time lapse movie of four cells obtained with the 30'-PR experimental scheme (first cell is the one shown on Fig. 1C,D). These movies were rotated based on the direction of the force at the start of the pull and corrected for cellular *xyz* drift. A single plane of each resulting z-stack is shown. The SiR-DNA channel (gray) is band-passed (see *Methods*). The GFP-ferritin channel (green) is the raw fluorescence. Arrows indicate when the force is exerted. The time stamp is indicated relative to the start of force exertion. Play speed is ~1200×). Uncompressed TIFF version of all 8 movies analyzed for the 30'-PR scheme (Fig. 2 and Fig. S3), along with the non band-passed channels, are available (Table S2).

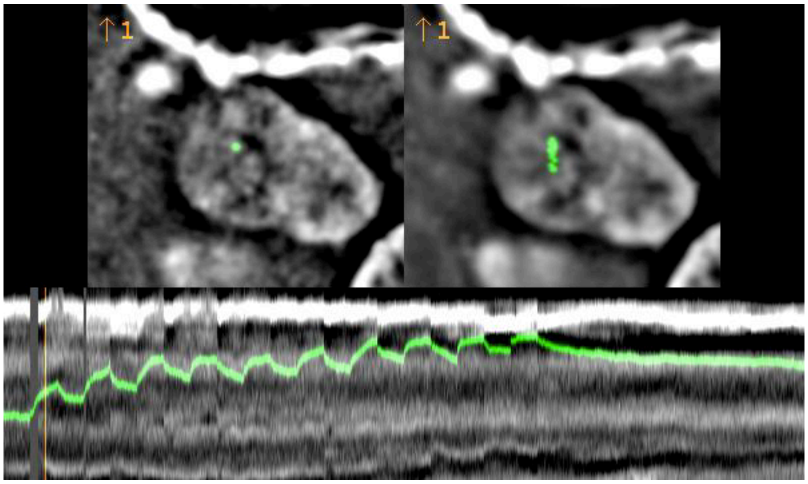


Movie S2. **Repeated 100 sec pull-release experiment.** Movie of the cell shown on Fig. 1E,F and analyzed on , Fig. S7 and Fig. S8 (100"-PR scheme). This movie was made with a fast frame rate (Δt=5sec) and as a single-plane acquisition. (**Top left**) The movie was rotated based on the direction of the force at the start of the pull and corrected for cellular *xy* drift. The gray channel (SiR-DNA) is band-passed and the green channel represents the center of mass of the locus (see *Methods*). Play speed is 100×. The magnetic micropillar is the black object with a bright contour at the top. The arrow indicates when the force is exerted. The number identifies the 10 individual pulls. (**Top right**) Time-projection images calculated over each 100 sec pull and release periods. (**Bottom**) Kymograph as shown on Fig. 1F, computed over the minimal *x*-range that contains the locus throughout the movie. The vertical bar identifies the frame shown on the top-left movie. Uncompressed TIFF version of this movie along with the non band-passed channels is available (Table S2).

Table S1 – **Amounts of MNPs and forces in all analyzed cells**

| Cell | Scheme | MNPs | | | Force (pN) | | |
|---|---|---|---|---|---|---|---|
| | | at the locus | total in the nucleus | % at the locus | start of pull | average of first 3 min | Max. over the pull(s) |
| 1 | 30’-PR | 1397 | 16240 | 8.6% | 0.308 | 0.310 | 0.625 |
| 2 | 30’-PR | 1218 | 45351 | 2.7% | 0.724 | 1.53 | 3.46 |
| 3 | 30’-PR | 1122 | 4754 | 23.6% | 0.084 | 0.084 | 0.090 |
| 4 | 30’-PR | 4467 | 15725 | 28.4% | 0.231 | 0.229 | 0.290 |
| 5 | 30’-PR | 816 | 2723 | 30.0% | 0.263 | 0.308 | 0.391 |
| 6 | 30’-PR | 350 | 3260 | 10.7% | 0.056 | 0.056 | 0.088 |
| 7 | 30’-PR | 2840 | 6106 | 46.5% | 0.410 | 0.423 | 0.431 |
| 8 | 30’-PR | 1003 | 1224 | 82.0% | 0.177 | 0.206 | 0.632 |
| 9 | 100”-PR | 1672 | 8642 | 19.3% | 0.735 | 0.530 | 4.26 |

Table S2 – **Summary of data and code availability**.

| | Description | Location |
|---|---|---|
| | **Centralized repository** for all the data and code described in this Table S2. | https://github.com/CoulonLab/Keizer-et-al |
| DATA | - **Registered and rotated TIFF files** (with raw and band-passed channels)<br> - All 8 cells analyzed for the 30’-PR experiment<br> - 100”-PR experiment, with time projections & kymograph (Movie S2)<br>- Data files with **trajectories and force time profiles** for all 9 analyzed cells<br>- **Fiji/ImageJ/Python scripts** and instructions used for generating these TIFF files and trajectory data files | https://zenodo.org/record/4674438 |
| DATA | - **Spinning-disk microscopy data for force calibration**. 6 pillars used in Fig. S5<br>- Force maps calculated on this data<br>- **Fiji/ImageJ scripts** and instructions used for generating these force maps | https://zenodo.org/record/4627062 |
| DATA | - **Microscopy data for single-MNP** intensity calibration<br>- **Fiji/ImageJ scripts** used for generating average single-MNP image | https://zenodo.org/record/4674531 |
| CODE | - **Python pipeline for concatenating** raw microscopy images | https://github.com/CoulonLab/chromag-pipeline<br>DOI: 10.5281/zenodo.4674417 |
| CODE | - MagSim, **Python library for magnetic simulations**<br>- Jupyter notebook for calibrating and generating maps in Fig. S4 & Fig. S5 | https://github.com/CoulonLab/MagSim<br>DOI: 10.5281/zenodo.4672595 |
| CODE | - **Python library for force inference** using different polymer models | https://github.com/SGrosse-Holz/rouselib<br>DOI: 10.5281/zenodo.4674399 |

### Theory for model-based analysis of locus trajectory

#### I. THEORETICAL SETUP

The biological system consists of a highly crosslinked, 4 Mb genomic locus embedded in human chr1, which itself contains 249 Mb. To model this system, we treat the chromosome as a bi-infinite continuous chain (i.e. $s \in \mathbb{R}$ below) and the locus as a compact ball attached to it at some point (labelled $s = 0$).

##### A. The chain

According to the Rouse model, the evolution of the chain conformation $x(s,t)$ follows the overdamped equation of motion

$$\gamma \dot{x}(s,t) = \kappa \partial_s^2 x(s,t) + \xi(s,t)\,, \tag{1}$$

where $\kappa$ encodes the stiffness of the chain, and the thermal kicks $\xi$ are a zero-mean Gaussian field with covariance

$$\langle \xi(s,t) \otimes \xi(s',t') \rangle = 2\gamma k_\mathrm{B} T \mathbb{1} \delta(s-s')\delta(t-t')\,. \tag{2}$$

The amplitude of the thermal noise is determined by the Einstein relation $D = k_\mathrm{B}T/\gamma$ for a free particle.

##### B. The locus

Since we model the chain as continuous, any single point along the conformation is infinitesimal in mass and extension. The only exception to this rule is the point $s = 0$, where we attach the compact ball that models the locus, i.e. we associate finite mass and volume with this point of the polymer conformation. In line with the overdamped nature of the motion we are studying, we will neglect the mass of this point right away. The finite extension means it will experience an additional drag force from the surrounding medium, slowing its motion at short time scales. More precisely, we would expect the locus to move diffusively at short time scales, where the effect of the rest of the polymer is still small. We do not see such a slowdown in the experimental MSDs (Fig. S6B), which shows that this viscous slowdown is negligible on the timescales we are studying. Following this argument, we also neglect the finite extension of the locus. Thus, finally, the locus is simply the point $s = 0$ along the chain, its finite size does not enter our model.

##### C. The force

A general force acting on the chain can be directly incorporated as an additional term $F(s,t)$ in the equation of motion (1). This can also be thought of as a non-zero mean of the thermal noise $\xi(s,t)$.

##### D. Solution

Taking all parts of our model together, we arrive at the equation

$$\gamma \dot{x}(s,t) = \kappa \partial_s^2 x(s,t) + \xi(s,t) + F(s,t)\,, \qquad \langle \xi(s,t) \otimes \xi(s',t') \rangle = 2\gamma k_\mathrm{B} T\, \mathbb{1} \delta(s-s')\delta(t-t')\,. \tag{3}$$

Noting that essentially this is a heat (or diffusion) equation, we can immediately write the solution as

$$x(s,t) = \int \mathrm{d}s' \int^t \mathrm{d}t' \, \frac{1}{\sqrt{4\pi\gamma\kappa(t-t')}} \exp\left( -\frac{\gamma(s-s')^2}{4\kappa(t-t')} \right) \left( \xi(s',t') + F(s',t') \right)\,. \tag{4}$$

This expression is the basis for all the following calculations. Especially, after some algebra one finds that the MSD of a point on the (force free) polymer is given by

$$\left\langle \left( x(s,t+\Delta t) - x(s,t) \right)^2 \right\rangle = \frac{2dk_\mathrm{B}T}{\sqrt{\pi\gamma\kappa}} \sqrt{\Delta t}\,, \tag{5}$$

where $d \equiv \mathrm{tr}\,\mathbb{1}$ is the number of spatial dimensions.

In the main text we occasionally write a heuristic powerlaw $\Gamma t^\nu$ for the MSD. From eq. (5) we identify $\Gamma \equiv \frac{2k_\mathrm{B}T}{\sqrt{\pi\gamma\kappa}}$ and $\nu = \frac{1}{2}$ for the Rouse model in one dimension.

#### II. FORCE INFERENCE

We are looking for a method to learn the applied force from observing the motion of the locus. From eq. (4) it is straight forward to calculate the reverse, the expected trajectory for a given force. We then proceed by inverting that relation, assuming that we know the position of the locus at discrete times.

Averaging over possible realizations of the thermal noise, we find the expected trajectory of the locus under an applied point force $F(s,t) = f(t)\delta(s)$:

$$\langle x(0,t)\rangle = \int^t \mathrm{d}t' \, \frac{f(t')}{\sqrt{4\pi\gamma\kappa(t-t')}} \,. \tag{6}$$

To invert this equation, we note that generally we only know the position of the locus at discrete time points $\{t_i\}_{i=1,\dots,N}$. Assuming the force to be constant during the intervals between these time points, $f(t) \equiv f_i \, \forall t_{i-1} \leq t < t_i$, we can evaluate the integral on these intervals and stitch the parts back together, finally yielding

$$x(t) = \frac{1}{\sqrt{\pi\gamma\kappa}} \Re \sum_{i=1}^{N} \left(\sqrt{t-t_{i-1}} - \sqrt{t-t_i}\right) f_i \,, \tag{7}$$

where we introduce the additional time point $t_0 < t_1$ and assume $f(t) = 0 \, \forall t < t_0$ (i.e. there is no force acting before the start of the observation). The expression $\Re\sqrt{x}$ is simply a neat way of keeping track of which terms contribute to the integral, in case $t < t_i$ for some $i$.[1] Now, evaluating the trajectory $x(t)$ at the discrete time points and writing $x(t_j) \equiv x_j$, we find

$$x_j = \frac{1}{\sqrt{\pi\gamma\kappa}} \Re \sum_{i=1}^{N} \left(\sqrt{t_j-t_{i-1}} - \sqrt{t_j-t_i}\right) f_i \tag{8}$$

or equivalently, writing $x \equiv (x_1, x_2, \dots, x_N)$ and $f \equiv (f_1, f_2, \dots, f_N)$:

$$x = \frac{1}{\sqrt{\pi\gamma\kappa}} M f \,, \qquad M = \begin{pmatrix} \sqrt{t_1} & & & \\ \sqrt{t_2} - \sqrt{t_2 - t_1} & \sqrt{t_2 - t_1} & & \\ \sqrt{t_3} - \sqrt{t_3 - t_1} & \sqrt{t_3 - t_1} - \sqrt{t_3 - t_2} & \sqrt{t_3 - t_2} & \\ \vdots & & & \ddots \end{pmatrix} . \tag{9}$$

For any given measured trajectory $\{(x_i, t_i)\}_{i=1,\dots,N}$ we can now assemble the matrix $M$ and then invert it, to finally find the forces acting on the locus between the sample points:

$$f = \sqrt{\pi\gamma\kappa} \, M^{-1} x \,. \tag{10}$$

*Calibration.* To use eq. (10) in practice, we have to know the prefactor $\sqrt{\pi\gamma\kappa}$. To that end we note that eq. (6) gives the response to a step force (i.e. switching on a constant force $F$ at $t = 0$) as

$$\langle x(t)\rangle = \frac{1}{\sqrt{\pi\gamma\kappa}} F\sqrt{t} \,. \tag{11}$$

We can therefore measure $(\pi\gamma\kappa)^{-1/2}$ as the slope of the force-displacement plot in Fig. 2C of the main text, which we find to be $0.231\,\mathrm{\mu m\,s^{-0.5}/pN}$. We use this value by default, unless explicitly stated otherwise on Fig. S8.

*Fluctuation-Dissipation.* Note that the prefactors of MSD (5) and response to a step force (11) differ exactly by a factor of $2dk_\mathrm{B}T$, as expected from the fluctuation-dissipation theorem (and the fact that MSD is a sum over $d$ independent spatial dimensions).

---

[1] $\Re$ signifies the real part, such that $\Re\sqrt{x} = \sqrt{x} \, \forall x \geq 0$ and $\Re\sqrt{x} = 0 \, \forall x \leq 0$. Since the boundary of the original integral is exactly such that the negative radicands will never appear, we can use this construction to incorporate that boundary, even though the sum runs over all the $t_i$ irrespective of where $t$ falls with respect to them.

#### III. COVARIANCES

The picture painted by eq. (10) is that of a bijective mapping between applied forces and observed trajectories. In reality, this mapping is blurred by thermal fluctuations, such that even a locus under the exact same applied force will move differently in repeated experiments. This in turn also implies that for a given observed trajectory, there is a multitude of reasonable force profiles. They will average to the one obtained by eq. (10), but in this section we want to study the possible deviations from this mean. To this end, we first study the variance in observed trajectories, if we were to resample the thermal noise. Propagating this through the inference will then tell us what force profiles are likely for the given trajectory at hand.

##### A. Covariance of locus trajectory

Starting from eq. (4), after some algebra one can show that

$$S(t,t') \equiv \langle [x(0,t)-x(0,0)] \otimes [x(0,t')-x(0,0)] \rangle = \frac{k_{\mathrm{B}}T}{\sqrt{\pi\gamma\kappa}} \mathbb{1} \left( \sqrt{t} + \sqrt{t'} - \sqrt{|t-t'|} \right) . \tag{12}$$

##### B. Error propagation

Moving to the discrete time picture of eq. (10), we can view $x \equiv (x(t_1), x(t_2), \dots, x(t_N))$ as a point in an $N$-dimmensional space. Similarly, $S$ from the previous paragraph becomes a $N \times N$ matrix with entries $S_{ij} \equiv S(t_i, t_j)$. Now, according to the Rouse model, the point $x$ is sampled from a Gaussian distribution with covariance matrix $S$. Consequently, we should view the inferred force $f$, which is a simple linear transform of $x$, as sampled from a Gaussian distribution with an accordingly transformed covariance matrix. Using eq. (10), this is

$$\mathrm{Cov}(f) = \pi\gamma\kappa M^{-1} \mathrm{Cov}(x) M^{-T} = \pi\gamma\kappa M^{-1} S M^{-T} , \tag{13}$$

where $M^{-T}$ is the inverse transpose of $M$.

#### IV. SUMMARY

Finally, we give a short summary of the inference method, given an observed trajectory $\{(x_i, t_i)\}_{i=1,\dots,N}$:

- calculate the matrices $M$ and $S$, using eqs. (9) and (12) respectively. These depend only on the sampling times and are in this sense "structural" quantities.
- use these structural quantities and the observed data to infer the whole ensemble of plausible force profiles. This is a Gaussian distribution with mean $\sqrt{\pi\gamma\kappa} M^{-1} x$ (eq. (9)) and covariance $\pi\gamma\kappa M^{-1} S M^{-T}$ (eq. (13)).
- this distribution specifies our full knowledge about the force acting on the locus. While the most interesting quantity is certainly simply the mean applied force profile, we can also explicitly sample from this distribution to obtain other examples of probable force trajectories.

The omnipresent prefactor $\sqrt{\pi\gamma\kappa}$ is calibrated from the force displacement profile shown in Fig. 2C of the main text.

#### V. VARIATIONS OF THE INFERENCE MODEL

Here we describe how the inference model changes upon variation of some of the underlying assumptions.

##### A. Finite tether

In the main inference scheme, we simply assume that the chromosome in which the locus is embedded stretches infinitely far in both directions. This seems to be a good approximation, as long as the whole chromosome indeed moves like a free polymer. This however does not necessarily have to be the case, as for example some parts of it might be fixed at the lamina (Lamina Associated Domains, LADs [1]), or at other points in the nucleus. For this reason, we ask how our inference scheme changes, if we assume the polymer on one side of the locus to have a finite extent, its end being fixed in space.

We still position the locus at $s = 0$, and assume the chain to extend to infinity for $s < 0$. In the positive direction, we add the boundary condition that $x(L,t) = 0\,\forall t$. Mathematically, we can explicitly take this boundary condition into account by modifying the fundamental solution according to the method of images. Equation (4) thus becomes

$$x(s,t) = \int_{-\infty}^{L} \mathrm{d}s' \int_0^t \mathrm{d}t' \frac{1}{\sqrt{4\pi\gamma\kappa(t-t')}} \left[\exp\left(-\frac{\gamma(s-s')^2}{4\kappa(t-t')}\right) - \exp\left(-\frac{\gamma(s-2L+s')^2}{4\kappa(t-t')}\right)\right] (\xi(s',t') + F(s',t')) \;, \quad (14)$$

which together with $\langle\xi\rangle = 0$ and $F(s,t) \equiv f(t)\delta(s)$ then gives the analog of eq. (6):

$$\langle x(0,t)\rangle = \int^t \mathrm{d}t' \frac{f(t')}{\sqrt{4\pi\gamma\kappa(t-t')}} \left[1 - \exp\left(-\frac{L^2}{\kappa(t-t')}\right)\right] . \quad (15)$$

Now following the same discretization scheme as before, we write

$$\begin{aligned} M_{ji} &= \Theta(t_j - t_i) \int_{t_{i-1}}^{t_i} \frac{\mathrm{d}t'}{2\sqrt{t_j - t'}} \left[1 - \exp\left(-\frac{L^2}{\kappa(t-t')}\right)\right] && (16)\\ &= \Theta(t_j - t_i) \left( \sqrt{t_j - t_{i-1}} \left[1 - \exp\left(-\frac{\pi^2\tau_L}{t_j - t_{i-1}}\right)\right] + \pi^{\frac{3}{2}}\sqrt{\tau_L}\,\mathrm{erfc}\left(\pi\sqrt{\frac{\tau_L}{t_j - t_{i-1}}}\right) \right., && (17)\\ &\qquad \left. - \sqrt{t_j - t_i} \left[1 - \exp\left(-\frac{\pi^2\tau_L}{t_j - t_i}\right)\right] + \pi^{\frac{3}{2}}\sqrt{\tau_L}\,\mathrm{erfc}\left(\pi\sqrt{\frac{\tau_L}{t_j - t_i}}\right) \right) \end{aligned}$$

where $\Theta$ is the Heaviside function (with the convention that $\Theta(0) = 1$) and we introduced $\tau_L \equiv \frac{L^2}{\pi^2\kappa}$ which is the Rouse time of a chain of length $L$. We also skipped the algebra involved in solving the integral. Note that for $\tau_L \to \infty$ this reproduces eq. (9). With this explicit expression for $M$ we can now run the inference scheme just like before.

##### B. Viscoelastic medium

The motion of certain proteins inside the nucleus has been observed to be sub-diffusive, with an anomalous exponent of $\alpha \approx 0.7$ [2]. It thus appears sensible to view the nucleoplasm as a viscoelastic medium. While this is certainly an effective picture, and this effective viscoelasticity might be due exactly to non-specific interactions with the surrounding chromatin, the motion of a Rouse polymer in viscoelastic solvent has been studied before [3]. One finds the generalizations of eqs. (5) and (6) as

$$\mathrm{MSD}(\Delta t) = \frac{2k_\mathrm{B}T}{\lambda} (\Delta t)^\nu \;, \quad (18)$$

$$\langle x(t)\rangle = \frac{\nu}{\lambda} \int_0^t \frac{f(\tau)\mathrm{d}\tau}{(t-\tau)^{1-\nu}} \;, \quad (19)$$

where $\lambda$ replaces our previous prefactor $\sqrt{\pi\gamma\kappa}$ and $\nu = \alpha/2$ generalizes the Rouse exponent of 0.5. We note that the presence or absence of a viscoelastic medium should be immediately apparent from the scaling behavior of the MSD. For our experiments this is shown in Fig. S6B and displays a clear square root law, from which we conclude that elastic effects in the solvent are negligible. For illustrative purposes, we nevertheless include a viscoelastic model with $\alpha = 0.7$ i.e. $\nu = 0.35$ in the model comparison in Fig. S8C. The force inference in this case works exactly as described above, with eq. (19) replacing eq. (6).

##### C. Dragging of surrounding chromatin

The Rouse model assumes no self-interaction of the chain, which is obviously a serious simplification. Here we will consider three ways that the pulled locus could be interacting with the surrounding chromatin, and how we can take that into account in the inference. Conceptual visualizations of the three models we will consider are shown in fig. 1.

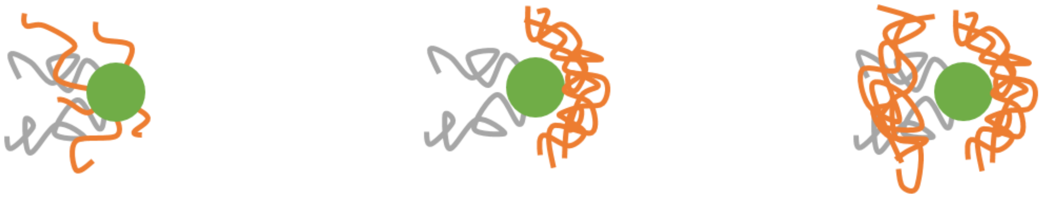

(a) Sticky chromatin (b) One-sided glove (c) Two-sided glove

FIG. 1: The three dragging models we will consider.

The idea behind the first type of models is that when moving, the locus has to push around the surrounding chromatin, which will accumulate in front of the locus as shown in fig. 1b. This is reminiscent of a baseball being caught in a catcher's glove, which is why we refer to these models as "glove models". In these models the locus is held back by the "glove" of accumulated chromatin in front of it, but is completely free to move backwards out of it (as it will for example upon recoil), in which case the glove will relax according to Rouse dynamics. This completely free recoil might appear reasonable, if we assume that the chromatin in the locus' path has already been pushed out of the way on the pull. If this is not the case, or the locus upon recoil takes a different path, we should expect a similar glove to build up behind the locus, as depicted in fig. 1c.

A different type of model, shown in fig. 1a, assumes that chromatin has strong non-specific interactions, meaning it will stick to the locus as it moves through the nucleus. In this case we will simply assume that at each point in time, some of the surrounding chromatin attaches to the locus and then moves with it.

The additional restoring forces generated by all of these models are then calculated using the same inference scheme as for the main locus itself. For every point $(x_i, t_i)$ in the given trajectory we generate a trajectory for a virtual particle that attaches to the locus at that point in time. Consequently, for $t < t_i$ this virtual particle is at rest at $x_i$, while for $t > t_i$ it follows the motion of the locus. How exactly this works depends on the model we use:

- for the sticky chromatin model, the trajectory for $t > t_i$ is simply exactly the one of the locus.[2]
- for the one-sided glove model, the virtual particle only stays attached as long as the locus is moving forward. As soon as it starts recoiling we calculate the relaxation of the virtual particle according to the Rouse model. This relaxation proceeds until the virtual particle comes back into contact with the locus, at which point the procedure starts again. This ensures that the virtual particle always stays ahead of the locus, and relaxes according to Rouse dynamics if it loses contact.
- the two-sided glove model works similarly to the one-sided one, except that particles that attach when the locus moves backwards will stay behind instead of ahead of it. Apart from that the procedure for obtaining their trajectories is exactly the same.

Examples of these virtual particle trajectories are shown in fig. 2.

Finally, we use the force inference on these trajectories to infer the additional restoring forces exerted by the dragged chromatin.

[1] B. van Steensel and A. S. Belmont, Cell **169**, 780 (2017).
[2] F. Höfling and T. Franosch, Reports on Progress in Physics **76**, 046602 (2013).
[3] S. C. Weber, J. A. Theriot, and A. J. Spakowitz, Physical Review E **82**, 011913 (2010).

[2] One could also imagine a finite lifetime for this stickiness, or a critical force that would break it, etc. For simplicity, we restrict ourselves to the basic model described.

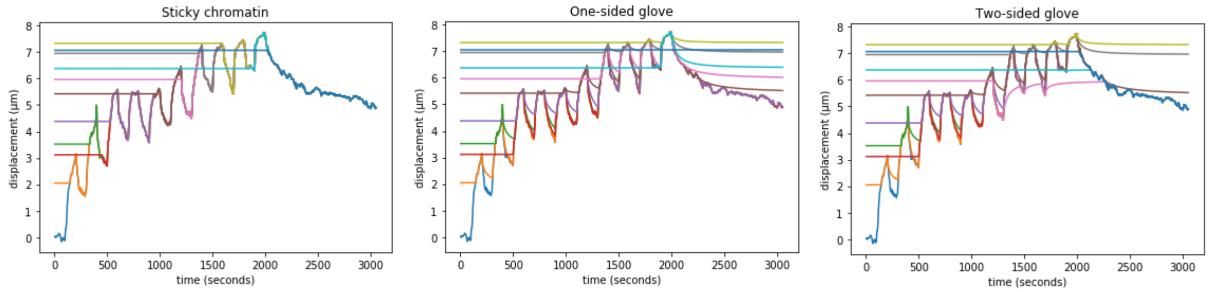


FIG. 2: Example trajectories for virtual particles used in the dragging models. For the sticky chromatin model, the virtual particles attach and then simply follow the motion of the locus. For the one-sided glove, we see the Rouse relaxation of the particles forming the glove when the locus recoils. The glove does not exert any force on the recoiling locus. In the two-sided glove model, there is a second glove behind the locus.
